# The Iron Metalloproteome of *Pseudomonas aeruginosa* Under Oxic and Anoxic Conditions

**DOI:** 10.1101/2025.01.15.633287

**Authors:** Mak A. Saito, Matthew R. McIlvin

## Abstract

*Pseudomonas aeruginosa* is a major contributor to human infections and is widely distributed in the environment. Its ability for growth under aerobic and anaerobic conditions provides adaptability to environmental changes and in confronting immune responses. We applied native 2-dimensional metalloproteomics to *P. aeruginosa* to examine how use of iron within the metallome responds to oxic and anoxic conditions. Analyses revealed four iron peaks comprised of metalloproteins with synergistic functions, including: 1) respiratory and metabolic enzymes, 2) oxidative stress response enzymes, 3) DNA synthesis and nitrogen assimilation enzymes, and 4) denitrification enzymes and related copper enzymes. Fe peaks were larger under anoxic conditions, consistent with increased iron demand due to anaerobic metabolism and with the denitrification peak absent under oxic conditions. Three ferritins co-eluted with the first and third iron peaks, localizing iron storage with these functions. Several enzymes were more abundant at low oxygen, including alkylhydroperoxide reductase C that deactivates organic radicals produced by denitrification, all three classes of ribonucleotide reductases (including monomers and oligomer forms), ferritin (increasing in ratio relative to bacterioferritin), and denitrification enzymes. Superoxide dismutase and homogentisate 1,2-dioxygenase were more abundant at high oxygen. Several Fe peaks contained iron metalloproteins that co-eluted earlier than their predicted size, implying additional protein-protein interactions and suggestive of cellular organization that contributes to iron prioritization in *Pseudomonas* with its large genome and flexible metabolism. This study characterized the iron metalloproteome of one of the more complex prokaryotic microorganisms, attributing enhanced iron use under anaerobic denitrifying metabolism to its specific metalloprotein constituents.

**Graphical Abstract:** 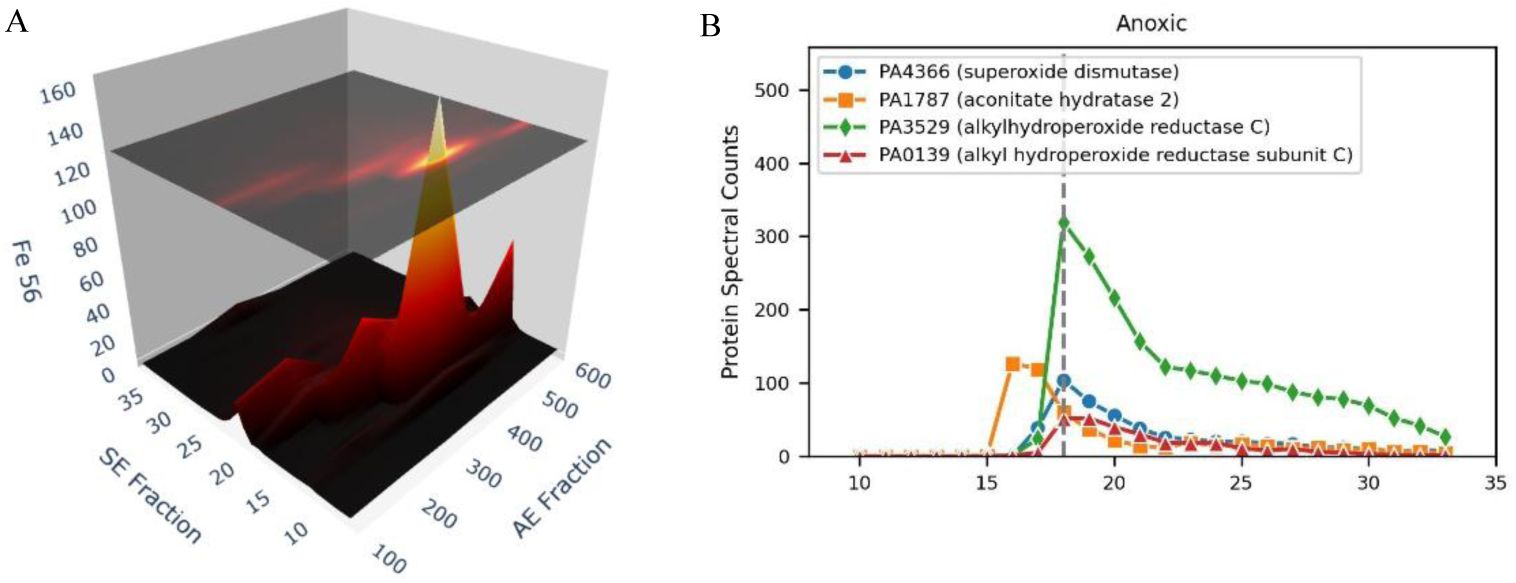

The iron metalloproteome of *Pseudomonas aeruginosa* was examined using native (non-denaturing) 2-dimensional chromatographic separation coupled to elemental and proteomic mass spectrometries. (A) Four major iron peaks were observed that corresponded to multi-protein complexes associated with respiratory, (B) antioxidant, DNA production, and denitrification functions, and associated iron storage and supply. The results suggest the presence of protein assemblies with potential roles in iron homeostasis and trafficking.

## Introduction

The pseudomonad bacterium are a group of widely distributed microbes found in both the environment and as pathogens [1,2]. The species *Pseudomonas aeruginosa* is a major contributor to infections in humans including in wounds and lungs of cystic fibrosis patients, as well as the cause of nosocomial infections such as pneumonia and septicemia [3]. Due to the prevalence of *P. aeruginosa* infections and increasing occurrence of antimicrobial resistance, the World Health Organization has classified it as a priority pathogen and recommended development of new antimicrobial strategies [4]. Furthermore, the role of metals is important in nutritional immunity, where immune cells target the availability of metals required by pathogens [5]. *P. aeruginosa* also occurs in natural environments, such as soils, waters, sediments and aquatic particles, as part of the microbial community that is decomposing organic matter [6–8].

Pseudomonads are capable of rapid growth, consistent with a copiotrophic lifestyle that allows them to proliferate quickly. Moreover, Pseudomonads are capable of multiple respiratory pathways that allows them to continue growth under suboxic or anoxic conditions, such as by growth on the oxidant nitrate via denitrification [9,10]. When infecting host tissues, *Pseudomonas aeruginosa’s* oxygen flexibility is advantageous, where it resists neutrophil cells’ oxidative bursts through the use of multiple superoxide dismutases, as well as also actively consuming oxygen to reduce the neutrophils’ effectiveness [11]. *Pseudomonas* has a large genome for a bacterium, with over 5000 genes in the core genome [12] and 5570 genes in the genome of PAO1 [13], reflecting these dynamic capabilities. Adaptations to acquire metals for nutrition is an important trait of *Pseudomonas aeruginosa,* and *P. aeruginosa* use of iron is particularly complex with regards to acquisition and homeostasis capabilities. As a result, an active area of development for antimicrobials involves drugs that target iron acquisition or trafficking, such antibiotic conjugated to siderophores, or inhibitors for iron release from ferritin [14–16]. Amongst *P. aeruginosa*’s sophisticated iron capabilities are the biosynthesis capability for two siderophores, pyoverdine and pyochelin, which are fluorescent and give cultures a characteristic visible coloration. *P. aeruginosa* also acquires iron through exogenous siderophores, through a ferrous iron transporter, and when complexed to heme, citrate, and catechols [17]. Metal insufficiencies caused through mutation of transporter systems often result in decreased pathogenicity, demonstrating the importance of metal nutrition in *P. aeruginosa* infections [18] and potential avenues for antibiotic treatment through inhibition of transporters and metal complexation by exogenous metallophores [14,17].

With this array of metal uses within *P. aeruginosa*, understanding the overall composition of this microbe’s metallome is of broad interest. The study of metalloproteins often focuses on characterization of individual proteins to provide essential mechanistic insights. In parallel, metallome studies can focus on the overall organismal or tissue metal content by bulk metal analyses and intracellularly localization of metals. Metalloproteomic techniques can complement these approaches by reconstructing the contribution of the composite of metalloproteins to the overall metallome. This approach combines native extraction of metalloproteins and their detection by dual mass spectrometry analyses to determine both metal and protein content simultaneously to assess major metalloproteins inventories within an organism [19–24]. In this study we conducted a metalloproteomic analysis of *P. aeruginosa* grown under aerobic and anaerobic conditions to assess how the iron metalloprotein inventory changes in response to these environmental conditions. A follow-up study will present the results of metals beyond iron in the *P. aeruginosa* metalloproteome.

## Methods

### Overview

This study combined the use of global, non-native detergent based extraction, proteomics to validate the oxic and anoxic experimental samples, prior to then further investigation of those samples by metalloproteomics, using native (no detergent) based extraction to maintain metal-protein coordination. The methods sections associated with each approach, as well as cultivation of biomass are described below.

### Isolation and Cultivation

Liquid media and plates for isolation were made with marine broth media base (Zobell’s media), which was prepared from 500 mL of 0.2 µm-filtered coastal seawater (Vineyard Sound) with the addition of 2.5 g peptone (Fisher Scientific) and 0.5 g yeast extract (BD Difco), with 7.5 g of agar (for plates only), followed by autoclaving for 40 min. Plates were streaked from 51 micron filter size fraction collected on a large volume submersible pump from 400m at Station 5 (0°S 202°E) on October 13, 2011 in the Central Pacific Ocean on the METZYME expedition (KM1128) aboard the R/V Kilo Moana and allowed to incubate at room temperature. Picked colonies from plates grown in liquid media displayed coloration characteristic of *Pseudomonas aeruginosa*. Once colonies had grown, colonies were restreaked and frozen at −80 in glycerol and shipped to the laboratory. Experiments were conducted with strain 2-54. While *P. aeruginosa* is known to be found in natural environments, including coastal environments, it is not often reported from open ocean (oceanic) environments. The sampling location was oceanic, although there were islands present in the broad geographic vicinity (Hawaii and Christmas Island of Kiribati). While this was an attempted isolation from the natural ocean environment, given genomic results and the lack of *Pseudomonas aeruginosa* observed in co-collected metagenomes and metaproteomes datasets [25,26], we infer that this isolate was likely a contaminant from the ship or its personnel during the isolation process.

### Genomic analyses

A bacterial isolate was genomic sequenced was sequenced at the Johns Hopkins Deep Sequencing and Microarray Core Facility using Oxford Nanopore and assembled using Canu, Nanopolish, and circulator pipeline. The genomes of isolates 1-54 and 2-54 were sequenced. 1-54 was assembled to a single circular chromosome of 6,455,702 base pairs, which had a 98.91% identity with the PAO1 genome, and was deposited to a repository (see Data Availability section). BLAST analyses and resulting annotations determined the microbe to be *Pseudomonas aeruginosa*. The genome of 2-54 did not assemble to a single closed circular genome, and was also found to be *Pseudomonas aeruginosa*.

### Genome Database for Proteomics Analysis

Given minor differences between PAO1 and our isolate, and the high-quality annotations associated with PAO1, the PAO1 genome was used for proteome annotation. PAO1 was downloaded from NCBI in January of 2019, allowing use of the well-established PA identification numbers.

### Anoxic Media

Anoxic media (anaerobic) treatments were conducted by autoclaving 1L bottles of media into followed by cooling, sterile-filtered sodium nitrate was added to a concentration of 0.88mM as an alternate oxidant to oxygen and enabling denitrification. The media was equilibrated with a loose cap in an anerobic glove box for several days to allow equilibration to the no oxygen environment. Anoxic bottles were then inoculated in the glove box, closed tightly, and vacuum heat sealed within a plastic bag. Oxic cultures were allowed to grow with a loose cap and moderate shaking to allow gas exchange. Cultures were grown in an incubator at 37°C with continuous moderate orbital mixing. Culture aliquots were harvested by centrifugation in 50mL centrifuge tubes at 3220 × *g* for 40 min at 4 °C using an Eppendorf 5810R centrifuge, the media was decanted, and pellets were frozen until analysis. 50mL aliquots of each treatment were extracted for both metalloproteomics and global proteomics.

### Protein analyses

Both global proteomics (using detergent based extraction) and native metalloproteomics (without detergent) were applied to the experiments conducted in this study, with the former applied to verify metabolic responses to environmental oxygen. Methods for both approaches are described below in their corresponding sections.

### Protein extraction for global proteomics

Pellets were resuspended in 5mL of protein extraction buffer (50mM HEPES pH 8.5 (Boston BioProducts #BB-2082), 1% SDS in HPLC grade water). All reagents in this protocol are made with HPLC grade water. Samples were heated at 95°C for 10 minutes and shaken at room temperature for 30 minutes. Samples were then spun for 30 minutes at 3220 x *g* in an Eppendorf 5810 centrifuge. Supernatant was removed from pellet and transferred to a Vivaspin 5K MWCO ultrafiltration unit (Sartorius Stedim #VS0611). Protein extract was concentrated to approximately 350 µL using the Vivaspin units, washed with 1mL of lysis buffer and transferred to a 2mL ethanol (EtOH) washed microtube (all tubes in global proteomics from this point on are EtOH washed). The Vivaspin units were then rinsed with small volumes of protein extraction buffer to remove all concentrated protein, and all samples were brought up to 400µL.

### Protein clean up and digestion

To purify (from detergent) and concentrate the global proteome protein extracts, a modified SP3 protein purification method was employed that uses magnetic beads to immobilize the proteins (Hughes et al., 2014). SpeedBead Magnetic Carboxylate Modified Particles (GE Healthcare #65152105050250 and #45152105050250) were prepared according to Hughes et al. (2014). 20 µL (20 µg/µL) of magnetic beads were added to 400 µL of extracted protein sample. Samples were heated at 37°C periodically to avoid precipitation. Samples were acidified to a pH of 2-3 by adding 50 µl of 10% formic acid. 2X volume (1100 µL) of acetonitrile was immediately added. Samples were incubated at 37°C for 15 minutes and then at room temperature for 30 minutes. Samples were placed on a magnetic rack, incubated for 2 minutes, supernatant was removed and discarded. Samples were washed 2 times removing and discarding supernatants with 1400 µL of 70% ETOH for 30 seconds on the magnetic rack. 1400 µL of acetonitrile was added to each sample for 30 seconds on the magnetic rack. Supernatant was removed and discarded. Samples air dried for approximately 4 minutes until acetonitrile had just evaporated. Samples were removed from the magnetic rack and beads were reconstituted in 90 µL of 50 mM HEPES pH 8.0.

### Protein reduction and alkylation

50 units (2 µL) of benzonase nuclease (Novagen #70746-3) was added to each sample and incubated at 37°C for 30 minutes. Samples were reduced by adding 20 µL of 200 mM DTT (Fisher #BP172-5) in 50mM HEPES pH 8.5 at 45°C for 30 minutes. Samples were alkylated by adding 40 µL of 400mM iodoacetamide (Acros #122270050) in HEPES pH 8.5 for 30 minutes at 24°C, occasionally heating to 37°C to prevent precipitation. The reaction was quenched by adding 40 µL of 200 mM DTT in 50mM HEPES pH 8.5. Total protein was then quantified according to the method described below (Total protein analysis).

### Peptide recovery and preparation

Acetonitrile was added to digested peptides at a concentration of ≥ 95% and incubated for 20 minutes at room temperature. Samples were then placed on the magnetic rack for 2 minutes and the supernatant was removed and discarded. 1400 µL of acetonitrile was added to samples on the magnetic rack for 15 seconds. Supernatant was removed and discarded. Samples were air dried for approximately 4 minutes, until acetonitrile had just evaporated. Beads were reconstituted in 90 µL of 2% DMSO and incubated off the rack at room temperature for ≥ 15 minutes. Samples were centrifuged slowly and briefly at 900 rcf to remove liquid from the tube walls. Samples were incubated on the magnetic rack for 15 minutes and supernatant containing peptides was transferred to a new ETOH washed 1.5 mL microtube. This step was repeated to ensure removal of all magnetic beads. 1% trifluoroacetic acid was added to samples for a final concentration of 0.1%. Samples were zip tipped with Pierce C18 tips (Fisher #87784) according to manufacturer’s protocol with a final resuspension in 25 µL of 70% acetonitrile, 0.1% formic acid. Samples were evaporated to approximately 10 µL in a DNA110 Speedvac (ThermoSavant). Samples were finally resuspended to a peptide concentration of 1 µg/µL in buffer B (2% acetonitrile, 01% formic acid).

### Total protein analysis

Total protein was quantified protein using 2µL of sample in duplicate and the BCA method (Thermo Scientific Micro BCA Protein Assay Kit #23235). Absorbance was measured on a Nanodrop ND-1000 spectrophotometer (Thermo Scientific). Standard curves were generated using albumin standard (Thermo Scientific #23210). The samples were then digested with trypsin (Promega #V5280) dissolved in HEPES pH 8.0 at a concentration of 0.5 µg/µL was added to samples at a 1:25 trypsin to protein ratio and incubated at 37°C overnight.

### Peptide analysis

Protein extracts were analyzed by liquid chromatography-mass spectrometry (LC-MS) (Michrom Advance HPLC coupled to a Thermo Scientific Fusion Orbitrap mass spectrometer with a Thermo Flex source). 1 µg of each sample (measured before trypsin digestion) was concentrated onto a trap column (0.2 x 10 mm ID, 5 µm particle size, 120 Å pore size, C18 Reprosil-Gold, Dr. Maisch GmbH) and rinsed with 100 µL 0.1% formic acid, 2% acetonitrile (ACN), 97.9% water before gradient elution through a reverse phase C18 column (0.1 x 250 mm ID, 3 µm particle size, 120 Å pore size, C18 Reprosil-Gold, Dr. Maisch GmbH) at a flow rate of 500 nL/min. The chromatography consisted of a nonlinear 160 min gradient from 5% to 95% buffer B, where A was 0.1% formic acid in water and B was 0.1% formic acid in ACN (all solvents were Fisher Optima grade). The mass spectrometer was set to perform MS scans on the orbitrap (240000 resolution at 200 m/z) with a scan range of 380 m/z to 1580 m/z. MS/MS was performed on the ion trap using data-dependent settings (top speed, dynamic exclusion of 15 seconds, excluding unassigned and singly charged ions, and precursor mass tolerance of ±3ppm, with a maximum injection time of 150 ms).

### Proteomics informatics

Mass spectra were searched against *Pseudomonas aeruginosa* PA01 proteome downloaded from NCBI in January of 2019 using Proteome Discoverer 2.0 using the SEQUEST HT algorithm (Thermo) with a parent tolerance of 10 ppm and a fragment tolerance of 0.6 Da. Proteome Discoverer output files were then loaded in Scaffold (Proteome Software) with a protein threshold maximum of 1.0% false discovery rate.

### Metalloproteomic Method: 2D Native Separation and Dual Mass Spectrometric Analyses

#### Protein Separation

Cell pellets from 50 mL centrifuge tubes were thawed in an anaerobic chamber (Coy Laboratory Products Inc.) and suspended in 5 mL of 50 mM TRIS buffer (pH 8.8, all buffers Chelex treated to remove background metals). The solution was transferred to a 15 mL PET plastic centrifuge tube (Fisher 055391) before sonication on ice for 2 min (1 second on/off pulses with a 5 min stop after 1 min, Fisherbrand Model 120 sonic dismembrator). The sonicated pellet was diluted to a total volume of 30 mL in two 15 mL tubes and centrifuged in a gas tight bucket rotor for 60 min at 3220 × g at 5 °C (Eppendorf 5810R). The supernatant was loaded onto a GE HiTrap Q HP anion exchange column (1^st^ dimension, AE hereon; single 1mL column) with a peristaltic pump at 0.25 mL/min. The column was then attached to an Agilent 1100 series HPLC pump and eluted with a gradient of sodium chloride at 0.5 mL/min, where buffer A was 50 mM TRIS buffer (pH 8.8), and buffer B was 1 M NaCl in 50 mM Tris buffer (pH 8.8). The gradient was 2 min at buffer A, followed by a linear increase to 60% buffer B over 12 min, then another linear increase to 100% buffer B over 6 min. Eluent was collected with a Bio-Rad 2110 fraction collector in 2 mL microcentrifuge tubes at a rate of 1 fraction every 2 min and stored on ice. Anion exchange column fractions were concentrated using 5000 Da molecular weight cutoff Vivaspin columns from 500 µL to 100 µL before size exclusion separation. Concentrated samples were then injected into a Tosoh Bioscience TSKgel G3000SWxl size exclusion column (2^nd^ dimension, SE hereon; 30 cm × 7.8 mm, pore size 250 Å) with guard column attached column (TSKgel G2000SWxl-G4000SWxl guard column, 4 cm x 6 mm) to a Michrom Paradigm MS4 LC system with a CTC Analytics HTC Pal autosampler, and samples were eluted with a 0.5 mL/min isocratic gradient of 50 mM Tris buffer (pH 7.5, 50 mM sodium chloride). Fractions were collected in 1 min intervals into 1.2 mL 96 well plates. 250 µL of each well was transferred to another 96 well plate for metals analysis by liquid handling robot. Each SE fraction in the oxic and anoxic metalloproteome represents a volume of 0.5mL.

#### Calibration of Size Exclusion Column

Protein size standards were eluted on the size exclusion column, tryptic digested, and analyzed by data independent analysis. The BEH200 SEC Test Mix (Waters Corp.) was used for calibration, and 100μL was injected and separated by the SE column (Tosoh Bioscience TSKgel G3000SWxl) at 0.5 mL min^−1^ then trypsin digested. The peptides were analyzed on a Thermo Fusion coupled with a Neo Vanquish HPLC (25 min non-linear acetonitrile gradient). The Fusion performed MS1 scans in the Orbitrap with 60K resolution at 380 to 1280 m/z with a cycle time of 0.5 s, and MS2 scans in the ion trap at normal resolution. Four proteins within the standard were measured: thyroglobulin (666 kD, 0.6 mg/mL), IgG (150 kD, 0.4 mg/mL), BSA (66.4 kD, 1mg/mL, and Myoglobin (15 kD, 0.4 mg/mL) Peptide peak areas were calculated using Skyline (Skyline-daily 23.1), and the average of the three most abundant peptides were used to identify the retention time of each protein.

#### Protein analysis of metalloproteomic fractions

In preparation for proteomic analyses, separated proteins were reduced, alkylated, and tryptic digested according to the following procedures. 200 µL of each sample was combined with 10 µL acetonitrile and 15 µL of 10 mM dithiothreitol in 100 mM ammonium bicarbonate. The samples were incubated for 30 min at 70 °C while shaking at 450 rpm. After cooling to room temperature, 30 µL of 20 mM iodoacetamide in 100 mM ammonium bicarbonate was added to each well and incubated for 30 min in the dark. 10 μL of 0.03 μg/μL trypsin (Promega Gold) was added to each well and incubated overnight at 37 °C while shaking at 450 rpm. Trypsin digested size fractionated samples were analyzed using a Michrom Advance HPLC system with reverse phase chromatography coupled to a Thermo Scientific Q-Exactive Orbitrap mass spectrometer with a Michrom Advance CaptiveSpray source. Each sample was concentrated onto a trap column (0.2 x 10 mm ID, 5 µm particle size, 120 Å pore size, C18 Reprosil-Gold, Dr. Maisch GmbH) and rinsed with 100 µL 0.1% formic acid, 2% acetonitrile (ACN), 97.9% water before gradient elution through a reverse phase C18 column (0.1 x 150 mm ID, 3 µm particle size, 120 Å pore size, C18 Reprosil-Gold, Dr. Maisch GmbH) at a flow rate of 500 nL/min. The chromatography consisted of a nonlinear 50 min gradient from 5% to 95% buffer B, where A was 0.1% formic acid in water and B was 0.1% formic acid in ACN (all solvents were Fisher Optima grade). The mass spectrometer monitored MS1 scans from 380 m/z to 1580 m/z at 70K resolution. MS2 scans were performed at 15K resolution on the top 10 ions with an isolation window of 2.0 m/z and a 10 second exclusion time.

#### Metals analysis of metalloproteins

250 µL of each size fractionated sample was combined with 50 µL of 30% hydrogen peroxide (MilliporeSigma Supelco, trace metal grade) and digested overnight. Further digestion was conducted by adding 500 µL of 5% nitric acid containing 1 ppb Indium and digesting overnight again. Samples were analyzed on an ICAP-Q inductively coupled plasma-mass spectrometer (Thermo) with an SC-4 DX FAST autosampler (Elemental Scientific, Inc.). Samples were analyzed in KED mode with helium as the collision gas. The following metal isotopes were analyzed: Fe 56, P 31, Co 59, Zn 66, Ni60, Cu63, Al27, Ti48, V51, Cr52, Mn55, Fe57, As75, and Mo95. Metals were calibrated to standard curves prepared the same as the samples (96 well plates, identical reagents), which were run between each set of anion exchange fractions.

#### Comparability between Oxic and Anoxic Metalloproteomes

Efforts were made to make the results between oxic and anoxic metalloproteomes consistent to allow comparisons. Total protein yields from the 1^st^ dimension were measured by total protein assay (see Total Protein Analysis above) for both oxic and anoxic treatments (Figure S1). This was the protein material loaded into the 2^nd^ SE dimension. The total protein was relatively similar between oxic and anoxic treatments, ranging between 9% and 56% more in the anoxic treatment in all cases, with the smallest difference occurring at the peak protein fraction for both consistent with similar protein loadings between each treatment (300-400mM; Table S1 and Figure S1).

#### Data analysis

The metals and protein datasets had slightly different ranges of data coverage, as summarized in Table S1. The 2D data was visualized in 6-8 anion exchange (AE) fractions as the first dimension (100-200mM, 200-300mM, 300-400mM, 400-500mM, 500-600mM, 600-800mM, 800-1000mM, 1000-1000mM (where the final range is a continued elution at the highest salt concentration), where the last two ranges were only available in the oxic metals dataset) and 38 size exclusion (SEC) fractions (fractions 1-38) in the second dimension for a grid of 304 samples. Two initial AE fractions containing largely dead volume (0 and 0-100mM) were not analyzed for metals or proteins. AE values were plotted as the lower value of their AE ranges (100, 200, 300, 400, 500, 600, 800, 1000). Metals and proteins were analyzed from a subset of SEC fractions, 7-38 for metals and 10-35 for proteins, resulting in ∼312 proteomic analyses (156 per treatment) and ∼448 metal analyses (256 and 192 for oxic and anoxic, respectively). The four datasets were organized in a uniform metalloproteome data template with consistent formatting of dataframes including empty fields where no data was present (since the extent of metal and protein coverage varied) to allow import into the Python notebook.

#### Informatics Methods

Data analysis and visualization were conducted by importing CSV files (see Data Availability section) into a customized Metalloproteomic Viewer written in Python (Version 3.10.9 Anaconda package) using Matplotlib and Plotly packages to create visualizations, identify maxima, and conduct optimization fittings and implemented as a Jupyter notebook (Version 7.2.2). Code for the Metalloproteomic-Viewer is available on GitHub at (see Data Availability section). The notebook workflow consisted of identifying major metal peaks, identifying the proteins with maxima on or adjacent to the metal peak, identification of known metalloproteins within the list by searching using Copilot AI. For Fe Peak 1, the contributions of proteins to the iron peak in the SE dimension was estimated using a local search optimization algorithm (L-BFGS-B; or Broyden, Fletcher, Goldfarb, and Shannon optimization) and minimize function in Python).

## Results and Discussion

### Global Proteomics

Cultures grown under aerobic and anoxic conditions were screened using proteomics to verify acclimation to each treatment prior to metalloproteome analyses. These results are briefly summarized here, with a more detailed proteomic analysis of the influence of oxygen on *P. aeruginosa* to be presented in a future study. The proteomic results confirmed acclimation to the presence or absence of oxygen, with characteristic proteins showing changes in protein abundance in response to O_2_, including the denitrification pathway, which contain numerous metalloenzymes involving Fe, Cu and other metals (the nitrate, nitrite, nitrous, nitric oxide reductases; Figure 1). A number of other metal-related proteins also were more abundant under anoxic conditions including molybdopterin biosynthesis proteins, cobalt chelatase CobN from cobalamin biosynthesis pathway [27], and the CopR copper regulator [28]. Anaerobic molybdopterin biosynthesis is consistent with Mo use within nitrate reductase.

**Figure 1.**
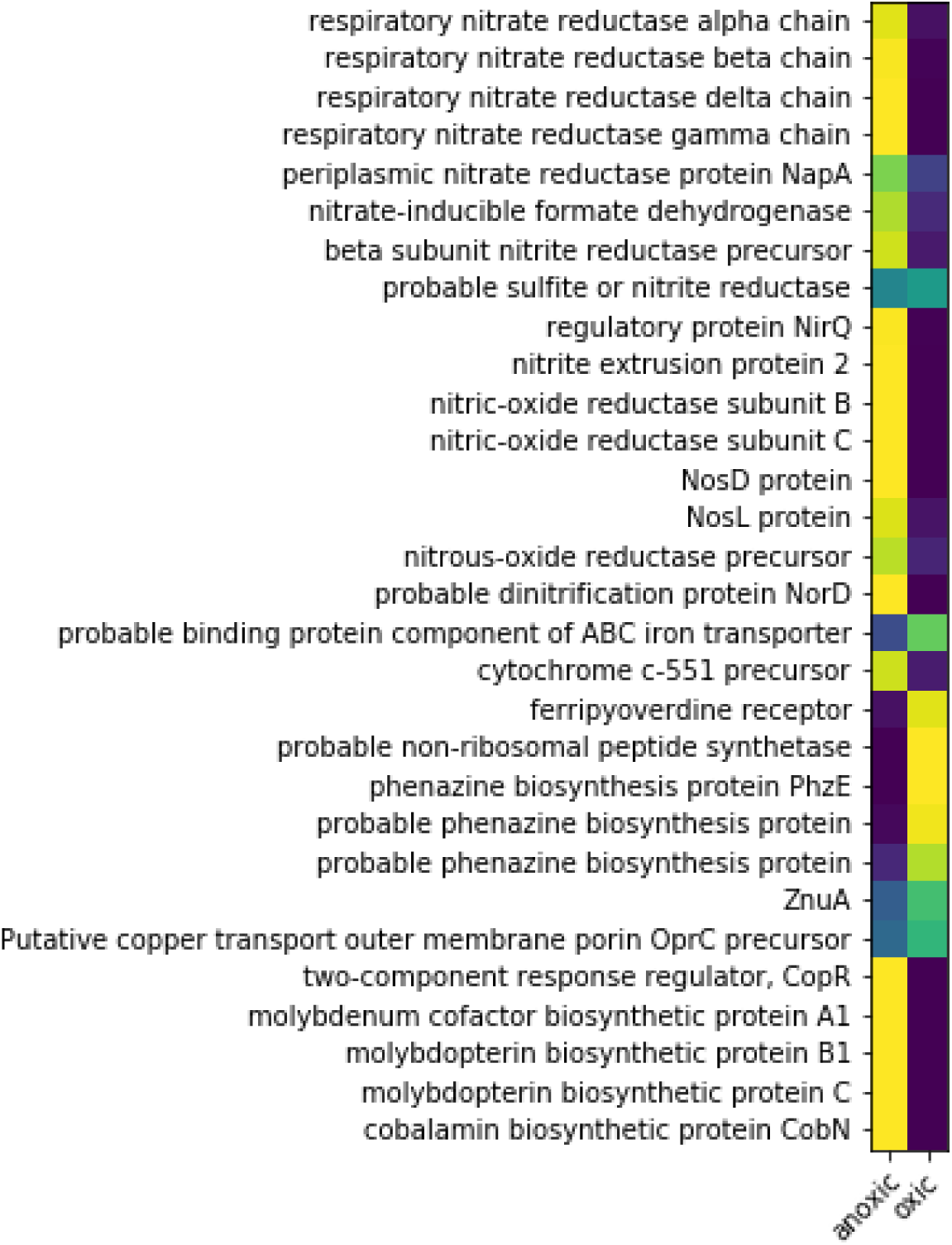
Global proteomic abundance data for selected metabolic and metal related systems in *P. aeruginosa* grown under aerobic and anaerobic conditions, where yellow is more abundant (values normalized to the sum of spectral counts for the two treatments; extraction triplicates were averaged). Strong responses were observed in nitrogen, iron, molybdenum and cobalt related systems. These global proteomes are sample splits of those used for the metalloproteomic dataset in this study.

The cobalt chelatase CobN, a component of the B_12_ biosynthetic pathway, increased in abundance under anoxic conditions. This contrasted a prior model that proposed aerobic B_12_ production, secretion and diffusion into anaerobic layers and uptake [29]. Because *P. aeruginosa* only has the oxygen-requiring enzymes for cobalt insertion in B_12_ biosynthesis (CobNST; as opposed to the oxygen independent CbiK/X found in other microbes), Crespo et al. proposed a model where B_12_ is produced aerobically in the upper biofilm layer, then must be secreted and diffuse to inner anaerobic layers where it is used for NrdJ (the B_12_-requiring for of the enzyme); and once depleted the Fe RNR-III (NrdD) is used below [29]. Ribonucleotide reductase enzymes also showed large changes in protein abundance between aerobic and anaerobic conditions (Figure 1; also see Fe Peak 3 section) [29]. Under aerobic conditions Class I NrdAB proteins were most abundant, with less of the B_12_-requiring class II NrdJab, and no detectable signal for the class III NrdD. Under anaerobic conditions, all three classes of RNR’s were present and in higher abundance than in oxic treatments (Figure 1). Similar to CobN, these results also contrast with the prior model, where a cascade of expression from NrdA to NrdJ to NrdG was expected to coincide with decreasing B_12_ availability [29]. Our experiments were conducted with mild orbital shaking, which discouraged accumulation of biofilms and may contribute to the differences with the prior study.

Overall, the results of the global proteomic survey demonstrated clear differences between oxic and anoxic treatments, particularly with regards to the denitrification apparatus. As a result, metalloproteomic analyses were conducted to determine changes in the metallome under oxic and anoxic conditions.

### Metalloproteomics Overview

Metalloproteomic analyses were conducted on sample splits from the oxic and anoxic treatments described above in the global proteomic results. This study focuses primarily on the iron metalloproteome, with the additional contextual Cu data. A future study will examine additional metals in the metalloproteome of *P. aeruginosa*.

The *P. aeruginosa* iron metalloproteome had four major peaks, labeled Fe Peaks 1-4 hereon, within the native 2-dimensional space created by the 1^st^ dimension of anion exchange (AE) and 2^nd^ dimension of size exclusion (SE) (Figures 2 and 3, Figure S2, Table 1). Notably the protein content of these Fe peaks was associated with major cellular functions, with Fe Peak 1 containing the oxic respiratory system and the bulk of iron storage. Fe Peak 2 was close in proximity to Fe Peak 1 and contained antioxidant functions. Fe Peak 3 was located higher in the AE dimension and contained enzymes associated with DNA synthesis and nitrogen use. Finally, Fe Peak 4 contained proteins associated with the denitrification capabilities and was found at the low end of the AE dimension and the high end of the SE dimension, consistent with more positively charged and smaller sized proteins, respectively. Notably Fe Peaks 3 and 4 showed increased Fe and protein content under the anoxic condition. The following sections describe the contents of the four major iron peaks, followed by a section about iron storage capabilities. A list of metalloproteins within each Fe Peak is provided in Table 2.

**Figure 2.**
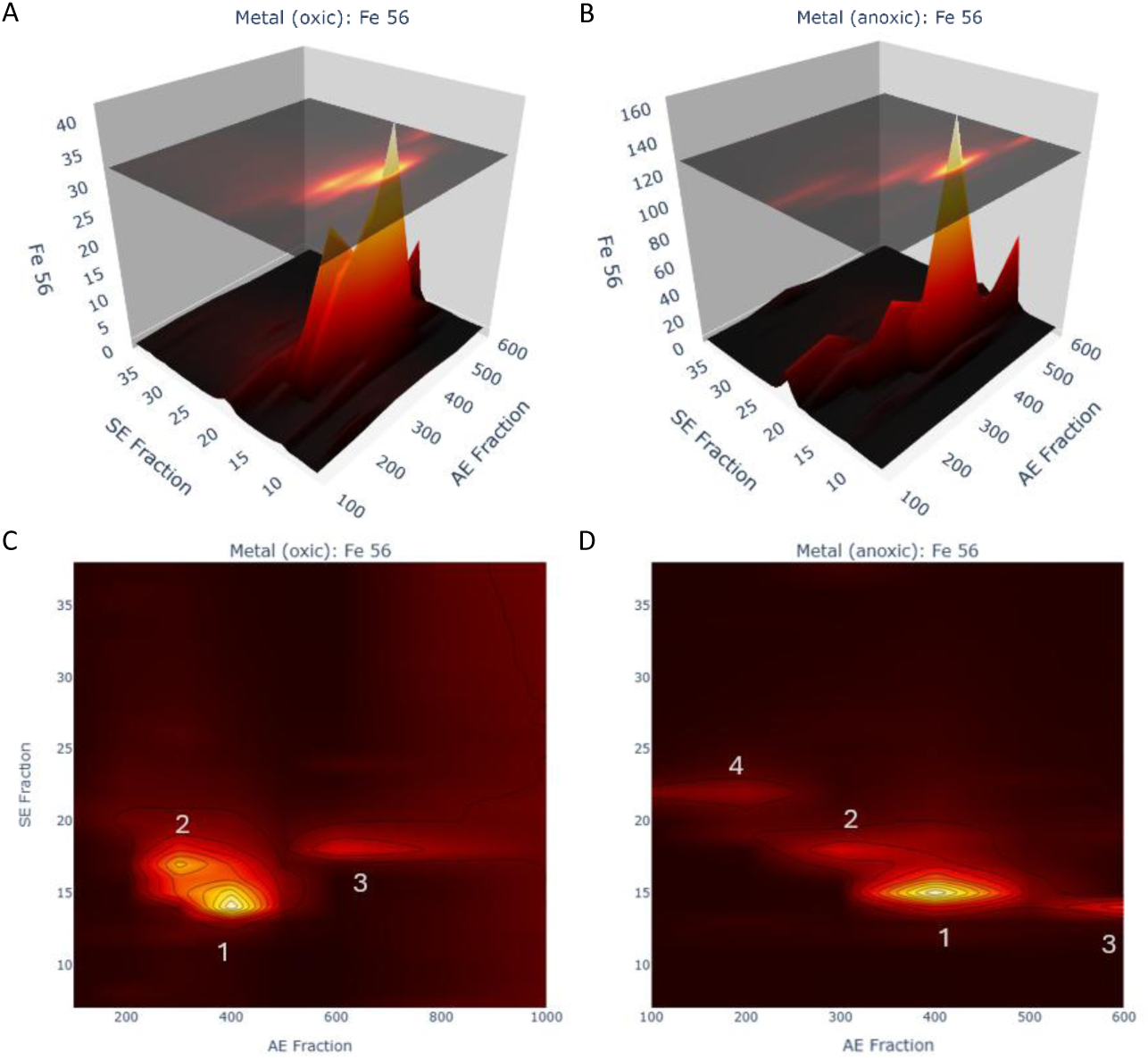
The Fe metalloproteome of *P. aeruginosa* under oxic (A and C) and anoxic (B and D) conditions in 3D and 2D reveals four major iron peaks (numbered, also see Table 1). The four peaks were associated with major cellular processes, 1) oxic respiration, 2) ROS detoxification, with 3) DNA production (dNTPs), and 4) denitrification. AE refers to the first dimension of anion exchange (in units of mM; note scale of oxic 2D is extended to 1000mM). SE refers to the second dimension of size exclusion and represents fraction numbers. Iron (Fe 56) is in units of ppb.

**Figure 3.**
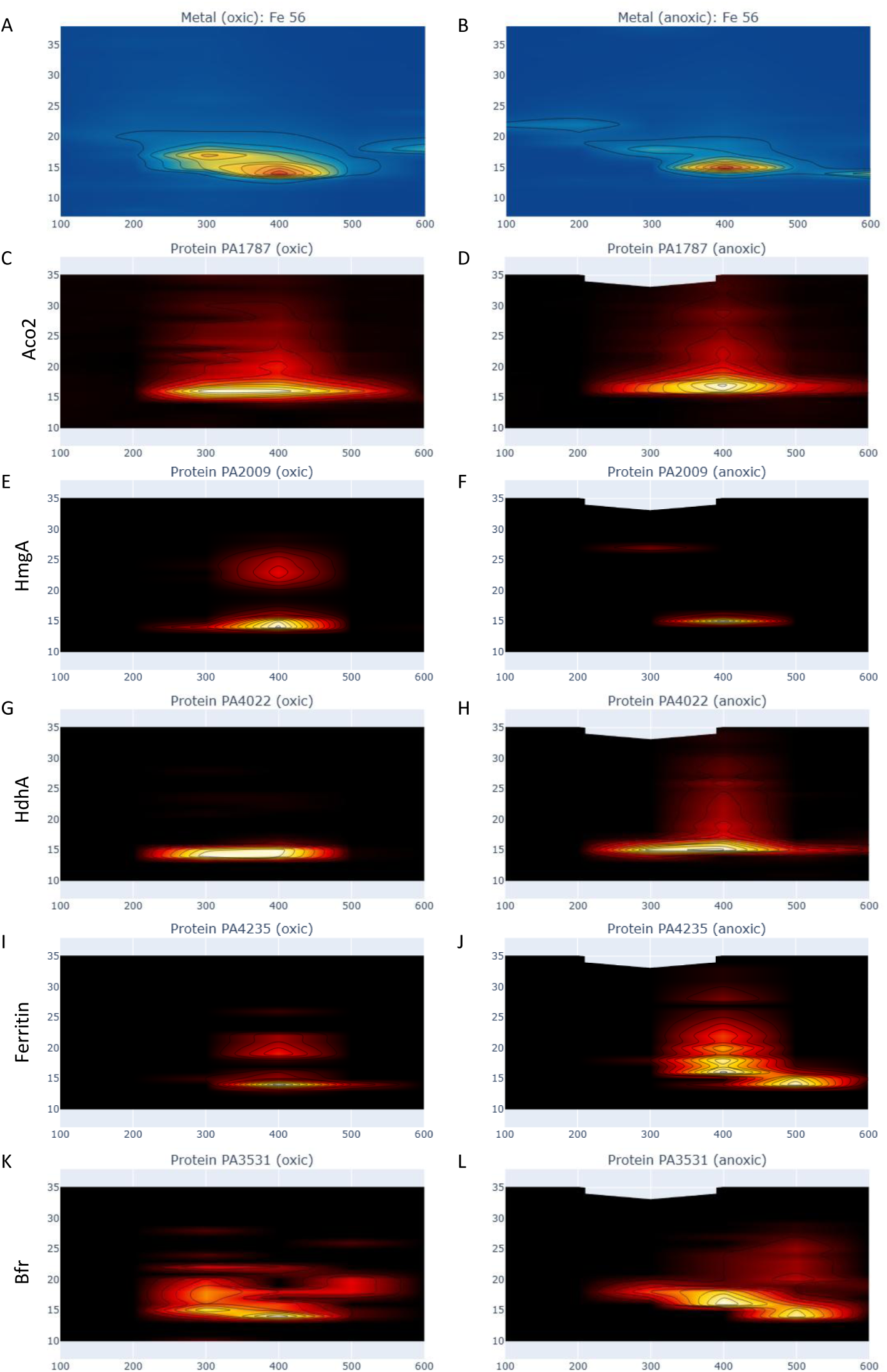
The iron metallome in 2D native space under oxic (A) and anoxic (B) conditions (similar to Figure 2 but with uniform scale for comparison to proteins), compared with the iron metalloproteins: aconitate hydratase 2 (C, D; Aco2, PA1787), homogentisate 1,2-dioxygenase (E, D; HmgA, PA2009), hydrazone dehydrogenase (G, H; HdhA, PA4022), bacterial ferritin (I, J; PA4235), and bacterioferritin (K, L; Bfr, PA3531). Note some iron proteins have broad peaks (PA1787, PA4022) or double peaks (PA2009, PA4235) that extend beyond the iron maximum region, which may be due to apo (unmetalled) or partially metalated isoforms.

**Table 1.** Major Fe peaks in the oxic and anoxic metalloproteome of *Pseudomonas aeruginosa*.

| Peak Number and Rank | Location Oxic (AE-SE) | Fe value Oxic (ppb) | Location Anoxic (AE-SE) | Fe value Anoxic (ppb) | Biochemical Function(s) |
| --- | --- | --- | --- | --- | --- |
| 1 | 400-14 | 40.5 | 400-15 | 158.96 | Respiration |
| 2 | 300-17 | 25.7 | 300-18 | 46.2 | Oxidative Stress |
| 3 | 600-18 | 11.6 | 600-14 | 57.8 | DNA production (via dNTP synthesis) |
| 4 | 200-20 | 4.1 | 200-22 | 28.7 | Denitrification |

**Table 2.** Iron proteins found within Fe Peaks 1-4 of the *Pseudomonas aeruginosa* metalloproteome. Individual proteins are labelled with a letter suffix after the iron peak number.

| Fe Peak Number | Protein (genome annotation) | PA ID | Monomer MW (kDa), Oligomer Stoichiometry, Comments |
| --- | --- | --- | --- |
| 1a | aconitate hydratase 2 | PA1787 | 94 kDa, monomer |
| 1b | homogentisate 1,2-dioxygenase (HmgA) | PA2009 | 48 kDa, 6mer |
| 1c | hydrazone dehydrogenase (HdhA) | PA4022 | 55 kDa, Hetero 36-mer, 192 heme [87] |
| 1d | catalase | PA4236 | 56 kDa, tetramer |
| 1e | bacterial ferritin | PA4235 | 18 kDa, 24-mer [68] |
| 1f | bacterioferritin | PA3531 | 19 kDa, 24-mer [68] |
| 2a | iron superoxide dismutase (SodB) | PA4366 | 21 kDa, homodimer [88] |
| 2b | quinone oxidoreductase | PA0023 | 35 kDa, tetramer [89] |
| 3a | glutamine synthase | PA5119 | 52 kDa, 12-mer [90], Fe-S |
| 3b | catalytic component of class Ia ribonucleotide reductase (NrdA) | PA1156 | 107 kDa, $\alpha_2\beta_2$ (active), $\alpha_4\beta_4$(inactive) [83] |
| 3c | NrdB, tyrosyl radical-harboring component of class Ia ribonucleotide reductase | PA1155 | 47 kDa, $\alpha_2\beta_2$ (active), $\alpha_4\beta_4$(inactive) [83] |
| 3d | probable bacterioferritin | PA4880 | 20 kDa, 24-mer [68] |
| 3e | aconitate hydratase 1 | PA1562 | 99 kDa, monomer |
| 3f | NirS nitrite reductase precursor | PA0519 | 63 kDa, dimer, heme [91] |
| 3g | electron transfer flavoprotein alpha-subunit | PA2951 | 31 kDa, heterodimer (57 kDa), Fe-S [92] |
| 3h | DNA-binding protein from starved cells, Dps | PA0962 | 17 kDa, 12-mer |
| 3i | ferric uptake regulation protein | PA4764 | 15 kDa, tetramer [93], Fe-S, Zn |
| 3j | L-cysteine desulfurase (pyridoxal phosphate-dependent) | PA3814 | 45 kDa, Fe-S assembly, homodimer [94] |
| 3k | class III (anaerobic) ribonucleoside-triphosphate reductase subunit, NrdD | PA1920 | 76 kDa, homodimer [55] |
| 4a | NirS nitrite reductase precursor | PA0519 | 63 kDa, dimer, heme [91] anoxic/oxic |
| 4b | NapA periplasmic nitrate reductase protein | PA1174 | 93 kDa, monomer or heterodimer [95,96] (111 kDa), anoxic |
| 4c | NirN | PA0509 | 54 kDa, heme insertion to NirS, monomer [97], anoxic |
| 4d | probable iron-binding protein IscU | PA3813 | 14 kDa, metamorphic S and D multimers [98], anoxic/oxic |
| 4e | Heme-transport protein, PhuT | PA4708 | 31 kDa, monomer [99] anoxic |
| 4f | cytochrome c4 precursor | PA5490 | 21 kDa, monomer [100], anoxic/oxic |
| 4g | cytochrome c-551 precursor | PA0518 | 11 kDa, monomer [101,102], anoxic |
| 4h | cytochrome c551 peroxidase precursor | PA4587 | 37 kDa, monomer [101,102], anoxic/oxic |
| 4i | cytochrome c5 | PA5300 | 13 kDa, anoxic |
| 4j | cytochrome c-type protein NapB precursor | PA1173 | 18 kDa, heterodimer [95,96] (111 kDa), anoxic |
|  | *Selected Additional Metalloproteins of Interest* |  |  |
| 2 | alkylhydroperoxide reductase C | PA3529 | 22 kDa, Non-iron, >300 kDa MPC [49] |
| 2 | alkylhydroperoxide reductase C | PA0139 | 21 kDa Non-iron, >300 kDa MPC [49] |
| 4 | nitrous-oxide reductase precursor | PA3392 | 71k Da, homodimer [103], anoxic |
| 4 | azurin precursor | PA4922 | 16 kDa, homotetramer [104], anoxic/oxic |
| 4 | probable glutathione peroxidase | PA0838 | 17 kDa, homodimer, anoxic/oxic |

While the metalloproteomic methodology was developed with the intention of maintaining the native metal-protein relationship, the results presented here also support notion that the native extraction methodology preserves some multi-protein complexes (MPCs). As a result, the co-elution of iron and other proteins within the some of the iron peaks described below may be neither coincidental nor due to limited chromatographic resolution, but remnants of *in vivo* protein-protein interactions. The final section of this manuscript briefly discusses this potential useful aspect of the metalloproteomic method.

#### Fe Peak 1

The largest peak (Fe Peak 1) was located at AE-SE location 400-14 in oxic conditions and 400-15 in anoxic conditions (Figure 2, Table 1). Six iron proteins co-eluted within this region (Figure 4, labeled 1a-1f below), aconitase (annotated as aconitate hydratase 2 PA1787), homogentisate 1,2-dioxygenase (HmgA, PA2009): and hydrazone dehydrogenase (HdhA, PA4022), catalase (PA02360), and two ferritins (annotated as bacterial ferritin PA4235 and bacterioferritin PA3531).

**Figure 4.**
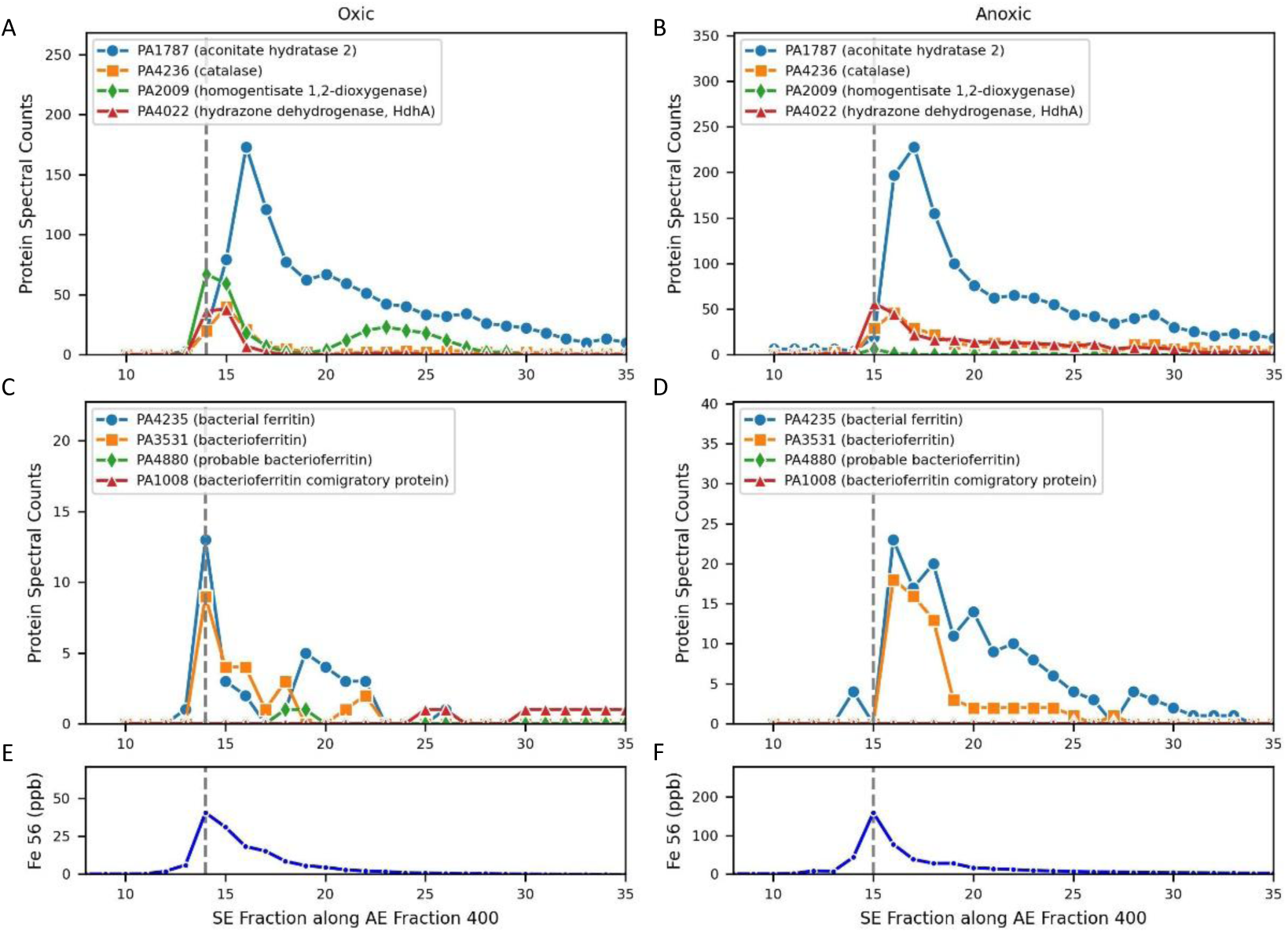
Iron proteins co-eluting with Peak 1 along the SE gradient within the 400 AE 1^st^ dimension under oxic (A, E) and anoxic (B, F) conditions. Note that all plots share the same x-axis. The iron maxima in this dimension aligns with the peak maxima of homogentisate 1,2-dioxygenase (PA2009) in both oxic and anoxic conditions, while the much larger aconitate hydratase 2 (PA1787) peak is offset by slightly offset by two SE fractions under both conditions, however this protein has a large broad distribution in native space (see Figure 2). The iron enzyme hydrazone dehydrogenase HdhA (PA4022) also co-eluted with Peak 1, and transitions from being a minor peak in the oxic conditions to a major peak in the anoxic conditions. (C-D) The ferritins (PA3531 and PA4235) align with the iron maxima under oxic conditions but are also slightly offset by SE 1 fraction under anoxic conditions. The iron peak has a shoulder in the 16-21 SE fraction range 9 (E-F), in both oxic and anoxic conditions from the confluence of these metalloprotein peaks.

*Fe Peak 1a*: The key respiratory protein aconitase was the most abundant iron protein with a large and very broad peak in both oxic and anoxic datasets that extended through multiple anion exchange fractions (AE 300-500 along SE 16 in oxic and AE 300-600 along SE 16-17; Figures 3C, 3D, 4A, 4B), reaching 173 and 228 spectral counts at each AE 400 maxima, respectively. However, both aconitase maxima were offset from the Fe Peak 1 maxima (along AE 400): under oxic conditions aconitase was at SE 16 versus Fe Peak 1 at SE 14, while under anoxic conditions aconitase was at SE 17 versus Fe Peak 1 at SE 15. There is a shoulder in Fe Peak 1 towards higher SE fractions, and the aconitase offset likely contributes to this. A similar offset of aconitase was observed in the metalloproteome of the marine bacterium *Pseudoalteromonas*, where a large second peak that eluted at a larger size fraction was offset from one that aligned with iron, and was interpreted as a potential apo form [21]. Given the breadth of aconitase elution in the *P. aeruginosa* metalloproteome, it also seems feasible that the metalated form occurs at the lower SE range, where the bacterioferritin and ferritin co-eluted (Figure 4C and 4D). The breadth of this peak could also reflect the extent of protein-protein interactions and resulting MPCs, where ferritin proteins can provide the source of iron to aconitase and other metalloproteins in Fe Peak 1 (see last section).

*Fe Peak 1b:* Another enzyme within Fe Peak 1, homogentisate 1,2-dioxygenase (PA2009, HmgA), was over 11-fold more abundant in the oxic treatment relative to the anoxic treatment (maximum of 67 and 6 spectral counts, respectively). This enzyme is involved in the degradation of aromatic rings including tyrosine and phenylalanine to produce the metabolite homogentisate (2,5-dihydroxyphenylacetate, HG), and requires oxygen for activity. It is a homo-6mer with an iron cupin site, where the cupin β-strands have a characteristic metal binding motif [30]. HmgA was observed to be one of the few genes downregulated in the hypervirulent Australian *P. aeruginosa* clinical strain AES-1 under chronic infection cystic fibrosis patients, relative to acute infection systems, and hence its regulation has been described as an important component of *P. aeruginosa’s* adaptation to the cystic fibrosis lung environment [31]. HmgA is also important in humans: a mutation to HmgA causes the rare genetic disease alkaptonuria, which causes an accumulation of homogentisic acid, due to the inability to break down tyrosine and phenylalanine, which polymerizes inside the body and as a dark pigmentation and causing severe arthropathy [32]. HmgA’s abundance within the largest iron peak and responsiveness to oxygen in this study is consistent with these prior studies and implies that if HmgA in the AES-1 strain responds similarly to oxygen as observed here, then hypoxic conditions would be characteristic of the cystic fibrosis lung microbiome environment, and the decreased expression of HmgA under those conditions could enable a shift of iron to other, non-oxygen requiring functions (see below and Fe Peak 4). [33]

*Fe Peak 1c:* The iron enzyme hydrazone dehydrogenase HdhA (PA4022) also co-eluted with Peak 1, and transitions from being a minor peak in the oxic conditions to a major peak that also aligns with the iron maxima in the anoxic conditions. HdhA transforms hydrazones, molecules with a C=N-N moiety, to hydrazides and acids. HdhA uses NAD^+^ as the oxidant to for the oxidative degradation of hydrazone molecules which PAO can grow on as a carbon source [34]. Prior studies observed that PAO1 grown on the hydrazone adipic acid bis(ethylidene hydrazide) as its carbon source upregulated *hdhA*, and mutants of the gene had depressed growth [34]. It is unclear to us why HdhA would be more abundant under anoxic conditions, since NAD^+^, which is regenerated by oxygen, is used by both aconitase and HdhA, but perhaps due to increased availability of hydrazones under anoxic and denitrifying conditions.

Fe Peak 1 was four times larger in anoxic conditions compared to oxic conditions, despite only having 9% more total protein used in the metallome and proteome analyses (Table S1). The alignment of HdhA with Fe peak 1 and HdhA’s increased abundance under anoxic conditions implies it contributes to the larger anoxic Fe Peak 1.

Comparing this with the prior HmgA, both proteins co-eluted at Fe Peak 1, but one protein was more abundant under oxic conditions (HmgA) and the other under anoxic conditions (HdhA), implying a shifting influence of proteins contributing to Fe Peak 1 with the change in oxygen availability.

*Fe Peak 1d:* The Fe metalloprotein catalase (PA0236) also eluted within Fe Peak 1 in both oxic and anoxic conditions. This enzyme is responsible for the deactivation of hydrogen peroxide. With aconitase being highly sensitive to reactive oxygen species that cause degradation of its iron-sulfur clusters [35], the close proximity of these two enzymes within Fe Peak 1 would be beneficial.

*Fe Peaks 1e and 1f:* Two ferritin iron storage proteins contributed to Fe Peak 1, and a comprehensive discussion of ferritins in the *P. aeruginosa* metalloproteome will be discussed in a later section.

*Optimization of contributions to Fe Peak 1:* quantitative reconstruction of Fe Peak 1 was attempted using an optimization approach (see Methods) along the AE 400 1^st^ dimension for both oxic and anoxic conditions (Figure 5, Table 3). Notably, the double peak of homogentisate 1,2-dioxygenase (HmgA, PA2009), where the second peak may have been apo based on lack of iron detected there, results in it being penalized in the optimization. When the second HmgA peak was manually adjusted to have zero protein spectral counts prior to optimization to test this notion of it having an apo form, it became a significant component to Fe Peak 1 (Table 3; Figure S3).

**Figure 5.**
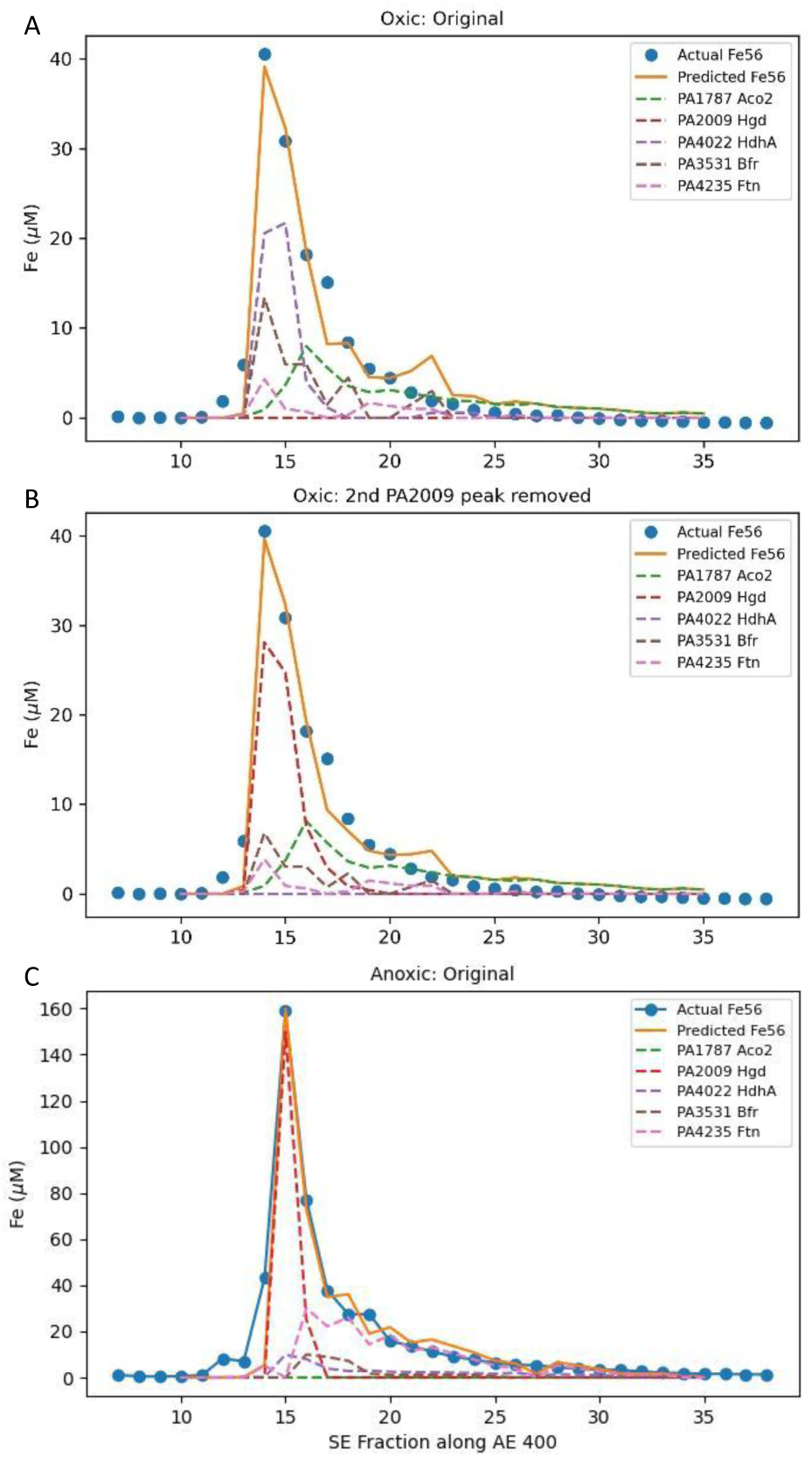
Optimization of protein contributions to Fe peak 1 under oxic (A) and anoxic (C) conditions. Note that PA2009 has a second peak at SE 23 under oxic conditions (see Figure 4) that may be an apo form. Because there is no iron peak at that location, the optimization results in a 0% contribution (A; Table 3). When the second peak of PA2009 was removed (B), PA2009 contributes 45.7% of the Peak 1 area and PA4022 goes to 0% (Table 3). In the anoxic treatment (C) the optimization produces a 0.0% (Table 3) contribution of PA1787, which similarly may be due to the broad and potentially apo forms of this abundant enzyme.

**Table 3.**
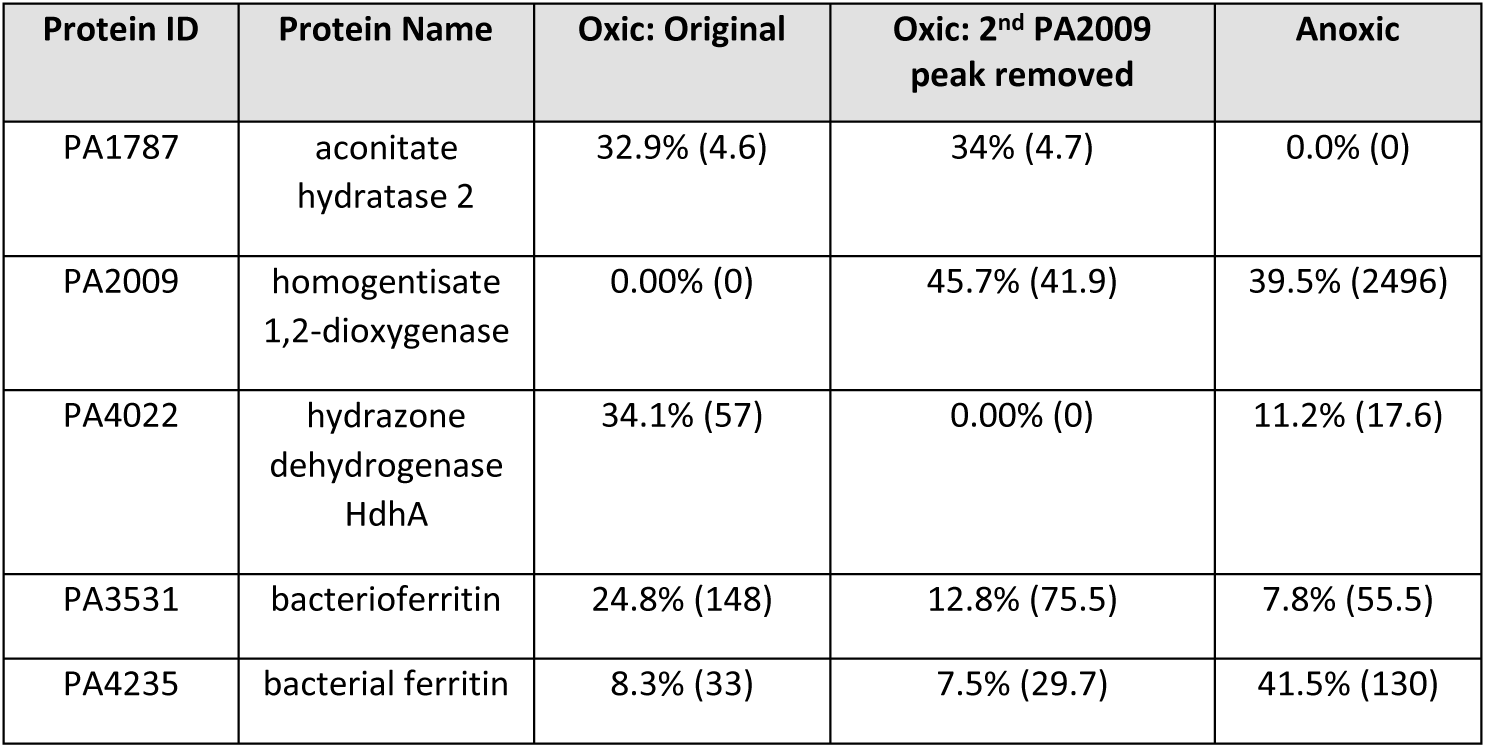
Optimization results for protein contributions to Fe Peak 1 (see Figure 5) as percent contributions to each optimization (oxic and anoxic). PA2009 (homogentisate 1,2-dioxygenase) had a second peak at higher SE fractions perhaps corresponding to an apo form, once removed PA2009 and PA4022 (hydrazone dehydrogenase HdhA) reversed their contribution estimates, illustrating the difficulty of determining the optimization when multiple proteins co-elute. Optimized coefficients are shown in parentheses multiplied by 100 for readability.

Given the multiple co-elution of iron metalloproteins within Fe Peak 1 (and other Fe Peaks, see below) in *P. aeruginosa*, the application of optimization approaches was challenging. Indeed, instead of independent metalloproteins that eluted uniquely based on their native charge and size properties as assumed in the optimization, there may be additional protein-protein associations creating co-dependencies between metalloproteins complicating assumptions of their independent co-elution. While this makes teasing apart the contribution of multiple iron proteins to each major iron peak difficult, there are potential biological implications of these co-elutions (see last section on multi-protein complexes).

*Summary of Fe Peak 1:* Fe Peak 1 appeared to be dominated by metalloproteins associated use of catabolism of carbon substrates and respiratory processes, such as via the Citric Acid cycle (aconitase), or degradation of aromatic rings (homogentisate 1,2-dioxygenase) under oxic conditions or hydrazones (hydrazone dehydrogenase) under anoxic conditions. In addition, catalase and iron storage proteins were both present, protecting Fe-S clusters (in aconitase) providing proximity to an iron source.

#### Fe Peak 2

The second largest iron peak within *Pseudomonas* was at 300-17 under oxic conditions and 300-18 (Figure 2, Table 1). Several iron proteins related to antioxidant activity had maxima associated with Fe Peak 2: the iron superoxide dismutase (SodB, PA4366), and quinone oxidoreductase (PA0023; Figure 6). Fe Peak 2 was also ‘diagonally’ adjacent to Fe Peak 1 described above and shared overlap of protein features: within oxic Fe peak 2 there was a shoulder towards the lower SE fractions, caused by a second peak at 300-15 that is likely associated with the aconitate hydratase 2 protein (PA1787; Fe Peak 1a) and its broad elution pattern mentioned above that continues at 300-16 (Figure 2, Figure 3A, 3E) and that contributes to both the AE 300 and AE 400 dimensions.

**Figure 6.**
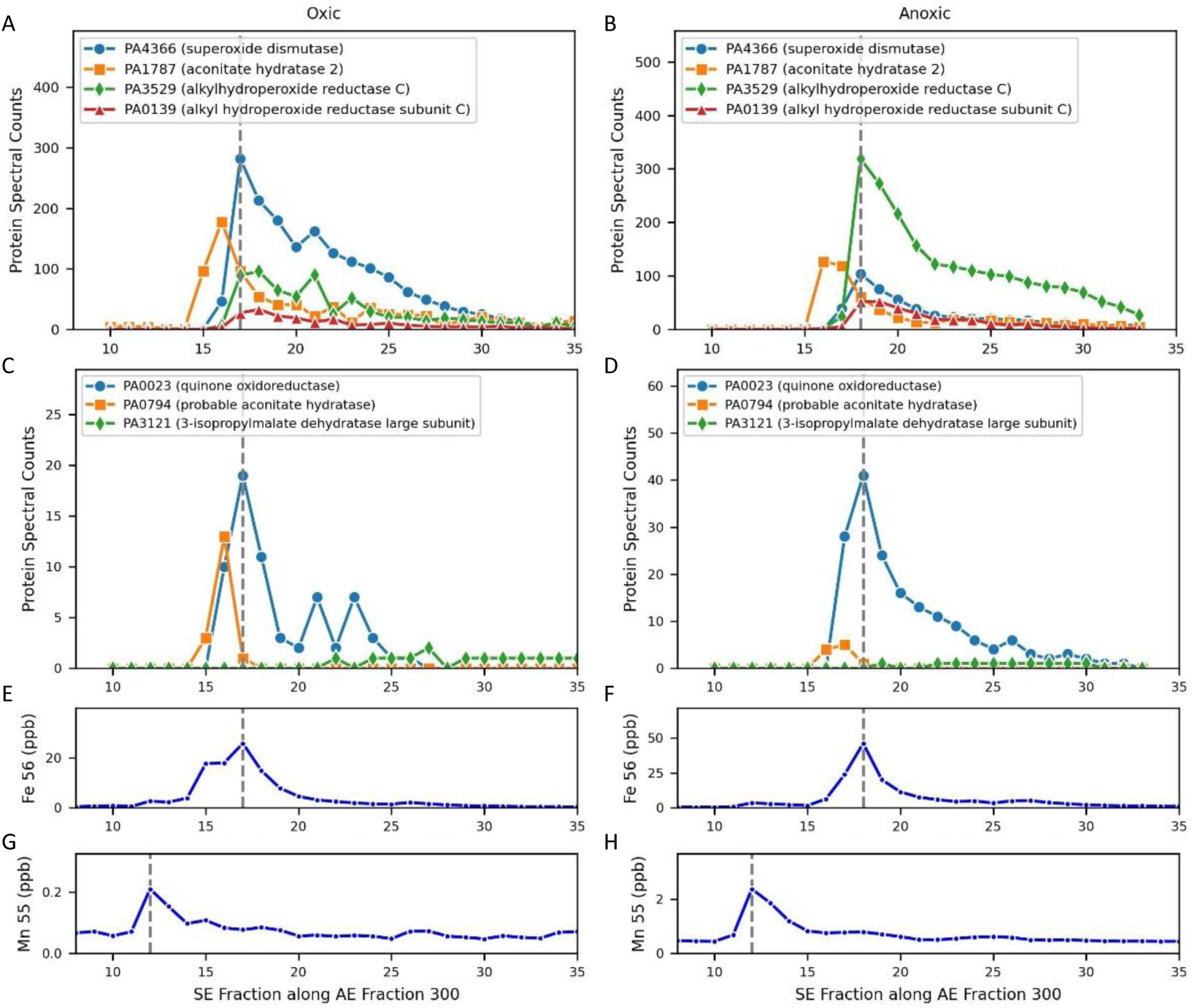
Iron proteins that contribute to Fe Peak 2, examined in 1D along the size exclusion fractions derived from the 300 AE dimension. The iron maxima in this dimension aligns with the peak maxima of superoxide disumutase (PA4366) in both oxic and anoxic conditions. Aconitate hydratase 2 (PA1787) is observed in Fe Peak 2, and it also observed in Fe Peak 1 (Figures 3, 4). As in Peak 1, it was also offset by 1-2 SE fractions under both conditions and has a large broad distribution in native space (see Figure 2). Two copies of the iron-regulated enzyme alkylhydroperoxide (PA3529 and PA0139) co-eluted with Peak 2, with PA3529 going from the 3^rd^ largest peak in oxic conditions to the largest peak in anoxic conditions, likely due to the need to address increased organic peroxides when growing on nitrate. and transitions from being a minor peak in the oxic conditions to a major peak in the anoxic conditions. Minor contributors to Fe Peak 2 included quinone oxidoreductase, a second copy of aconitate hydratase, and 3-isopropylmalate dehydratase large subunit (middle panels). There is a shoulder in Fe Peak 2 on the left side of the oxic peak at SE 15, likely contributed to by PA1787. The dotted grey line in all panels represents the local of the Fe maximum in AE 300, and the Fe anoxic peak was larger than under oxic conditions.

*Fe Peak 2a:* In *P. aeruginosa*, FeSOD (SodB, PA4366) has been reported to be produced constitutively, including under anaerobic conditions, while the MnSOD (SodA, PA 4468) is only produced under oxic stationary phase conditions [36,37]. These SODs, with catalase, contribute to defense against oxidative bursts from host neutrophil cells [11]. Our results are consistent with these prior findings, where the MnSOD was not identified in the oxic or anoxic metalloproteomes (see supplemental datasets within Data Availability section), which were both harvested in exponential growth, and the FeSOD was abundant in both treatments. Moreover, the Mn maximum was not aligned with the FeSOD implying no substitution in this SOD isoform (Figure 6G and 6H).

*Fe Peak 2b:* Quinone oxidoreductase (PA0023) is an antioxidant enzyme catalyzes the reduction of quinones preventing the formation of semiquinones that generate reactive oxygen species. [38] In bacteria the enzyme plays a critical role in the electron transport chain, catalyzing the transfer of electrons from NADH to quinone, and contains multiple iron-sulfur clusters [39]. Interestingly, NAD(P)H quinone oxidoreductase is often highly expressed in cancers, and hence is considered a drug target [40]. PA0023 was present in much lower abundance than SodB and aconitate hydratase 2, and hence was likely a minor contributor to Fe Peak 2.

*Fe Peak 2 Alkylhydroperoxide reductases*: Although not iron proteins, two versions of another antioxidant enzyme, alkylhydroperoxide reductase C (AphC, PA3529 and PA0139), also co-eluted at Peak 2 (Figures 6A and 6B). AhpC has been observed to be regulated by iron in *Campylobacter jejuni* [41]. The PA3529 and PA0139 sequences were checked using METATRYP software and do not share any tryptic peptides, meaning their proteomic identifications in the metalloproteome datasets here were independent of each other, and hence both proteins were present [42]. This enzyme likely has an important role in the physiology of *P. aeruginosa*, as observed in other organisms, for example AhpC is the primary scavenger of hydrogen peroxide in *E. coli* [43,44]. AhpC contributes to the cell’s antioxidant capability by removing organic peroxides such as peroxynitrite and is part of the 2-Cys peroxiredoxin family. As a result AhpC has been described as the “most widely distributed” reactive nitrogen intermediate (RNI) resistance gene [44]. Hare et al. [45] observed the PA3529 alkylhydroperoxide reductase C to have increased abundance under peroxide stress treatments in *P. aeruginosa* strain PAO1, in addition to positive responses from superoxide dismutase and catalase. PA3529 immediately precedes bacterioferritin-associated ferredoxin Bfd (PA3530) and bacterioferritin (PA353) in the genome, implying co-regulation of this gene neighborhood, as observed in prior transcriptome and proteome studies [46]. This is consistent with prior studies showing a dependence of catalase activity on bacterioferritin and susceptibility to hydrogen peroxide [47]. AhpC also exhibits chaperone activity, preventing misfolding and thermal aggregation of proteins [48]. This dual function of AhpC is thought to convey its adaptability to oxidative and environmental stresses [48]. AhpC exists as either a dimer or a decamer in physiological solutions, and has also been observed in multi-protein globules with molecular weights exceeding 300 kDa [49].

Both AhpC proteins were present in the native separation (metalloproteomic) analysis contributing to Fe Peak 2, with PA3529 having higher spectral counts than PA0139 in both oxic and anoxic conditions (Figure 6A, 6B). This trend was observed in the global proteome data, where PA3529 had 25% and PA0139 62% higher spectral counts under anoxic treatment relative to the oxic treatment. PA3529 went from being less abundant in spectral counts than superoxide dismutase under oxic conditions, to being more abundant than superoxide dismutase under anoxic conditions. The minor AhpC (PA0139) also increased under anoxic conditions from 33 to 52 spectral counts. Both AhpC proteins were minor constituents within the adjacent Fe Peak 1 (centered between SE 11-13 in both cases). Given the switch to nitrate-based respiration in the anoxic treatment (Figure 1), the increased abundances of both AhpC’s was likely induced by increased production of organic peroxides like peroxynitrite, and the need to deactivate them to prevent toxicity. This is consistent with Ahp in *P. aeruginosa* being regulated by the *ohr* system that senses organic peroxides, yet is unresponsive to hydrogen peroxide [50,51] and is consistent with both AhpC copies having increased abundance under anoxic denitrifying conditions, where organic peroxides can occur. Given the prevalence of hypoxic conditions within *P. aeruginosa* infections and its increased abundances, inhibitors to the AhpC could be a potential antimicrobial therapeutic target [52].

*Summary of Fe Peak 2:* Overall, Fe Peak 2 displayed key antioxidant capabilities (iron enzymes SodB and quinone oxidoreductase, and the non-iron containing AhpC) being deployed both under oxic and anoxic conditions (AhpC for reactive nitrogen intermediates) in close proximity (in native chromatographic space) to respiratory systems of Fe Peak 1 and iron storage systems.

#### Fe Peak 3

The third largest iron peak in the *P. aeruginosa* metalloproteome occurred in the 1^st^ dimension AE 600 mM and AE 500mM fractions. Here we consider oxic peak AE 600-SE 18 and anoxic peak AE 600-SE 14 both as Fe Peak 3 given their relative proximity in the iron visualizations and some shared proteins (Table 1, Figures 2, 3, 7A-F; note Figure 2 has expanded AE range to AE 1000 which includes Fe Peak 3). Notably the iron in Fe peak 3 was 4-5 fold larger in the anoxic treatment than the oxic treatment (Figure 7E and 7F for AE 600, and 7I and 7J for AE 500 dimensions). The total protein in the anoxic 600mM AE fraction was only 23.1% larger than the oxic (Table S1), not enough to account for the 5-fold Fe difference, hence we interpret this as a biological signal. After discussion of a technical issue, the identification of metalloproteins within oxic and anoxic treatments of Fe Peak 3 are briefly outlined (Table S2, also described as Fe Peak 3a-k), followed by discussion of selected proteins and their biological role across both treatments, and overall assessment of Fe Peak 3’s metabolic roles.

**Figure 7.**
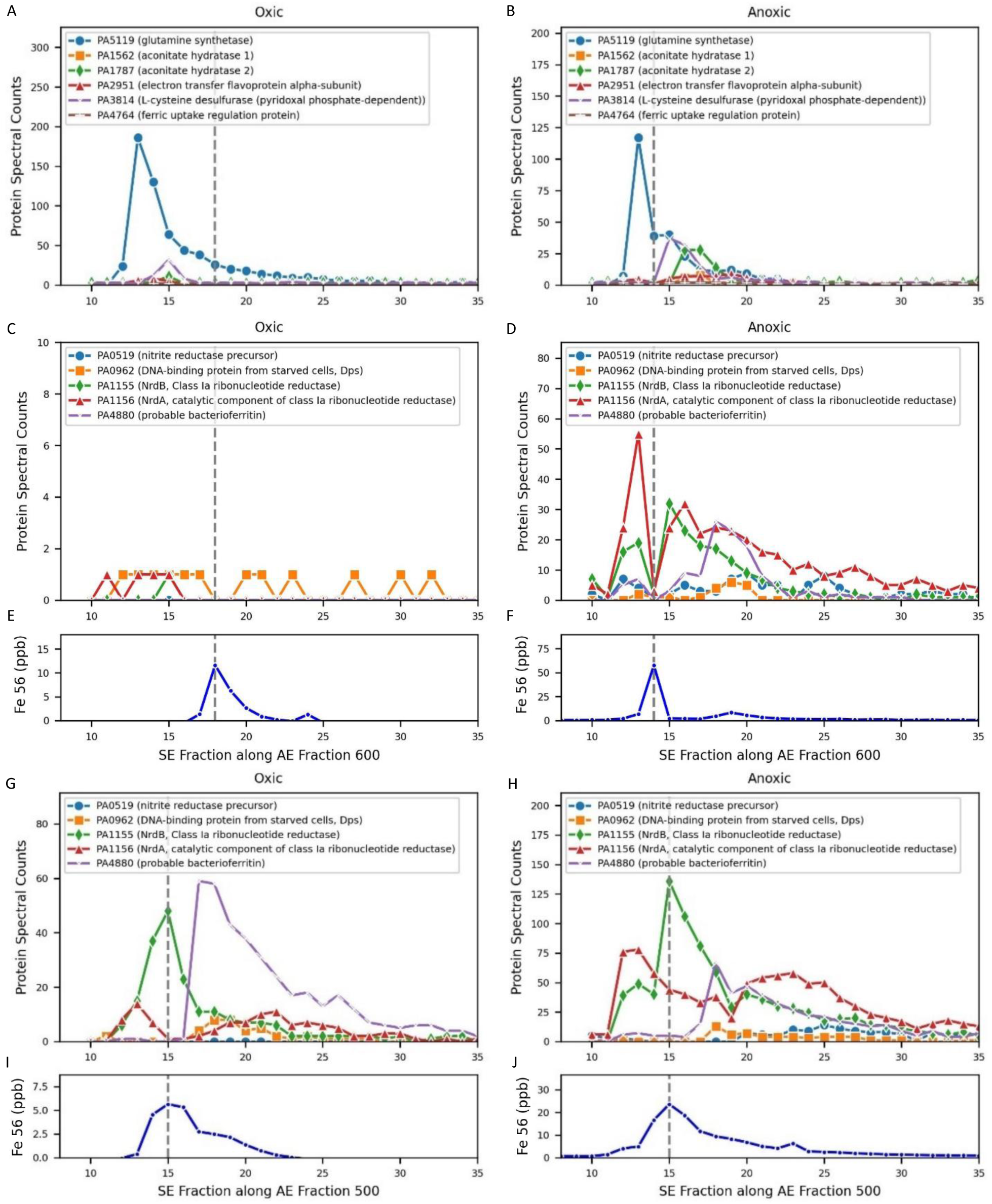
Proteins eluting at Fe Peak 3 in the AE 600 (A-F) and AE 500 (G-J) 1^st^ dimension fractions. Dotted grey represents the iron maxima.

There was a divot in overall signal intensity of protein abundances at anoxic Fe enzymes at AE600-SE14 (Figure 7B, D) where the iron maxima occurred. This divot was not consistent with protein data points on either side of its chromatographic elution, resulting in a ‘missing peaks’ for those proteins. The cause of the divot appears to be an instrument or sample processing issue rather than representing an actual biological signal, as evidenced by the overall mass spectrometry ion count signal in the sample being 1/10^th^ that of the surrounding samples in the raw data files (raw data available within supplemental raw datasets submitted to PRIDE). To provide redundancy of perspective for the missing peaks, we also examined for overlap of proteins and iron with the adjacent AE 500 dimension, where there was an Fe peak at SE 15 for both, relatively close to the anoxic AE600-SE14, and a bit more removed from the oxic AE600-SE18. As a result, AE 500mM is included in the Fe Peak 3 discussion here (Figure 7G-J, Table S1).

*Proteins in Oxic Treatment:* At oxic AE 600-18 there were no proteins with maxima that co-eluted at that precise location. Of all 33 proteins identified at that location, only three were known iron proteins (Figure 7A, C, E): glutamine synthase (PA5119), aconitate hydratase 2 (PA1787), and L-cysteine desulfurase (pyridoxal phosphate-dependent, PA3814). As mentioned above, the oxic Fe Peak 3 signal was much 5-fold smaller than the anoxic treatment.

*Proteins in Anoxic Treatment:* At anoxic 600-14, there were also no proteins with maxima at this precise location, although this was likely caused by the signal divot mentioned above. Of the 59 proteins detected at that location, the three iron proteins found in oxic AE 600-18 were present here, as well as additional the potential iron proteins (Figure 7B, E, F): aconitate hydratase 1 (PA1562), nitrite reductase precursor (PA0519), electron transfer flavoprotein alpha-subunit (PA2951), DNA-binding protein from starved cells, Dps (PA0962), and ferric uptake regulation protein (PA4764), NrdA, catalytic component of class Ia ribonucleotide reductase (PA1156) and NrdB, tyrosyl radical-harboring component of class Ia ribonucleotide reductase (PA1155), class III (anaerobic) ribonucleoside-triphosphate reductase subunit, NrdD (PA 1920). There were 8 unannotated proteins that eluted with maxima at 600-18 under anoxic conditions (PA3309, PA2765, PA0916, PA2817, PA3982, PA2151, PA3850, PA0457).

*Fe Peak 3a:* Glutamine synthetase (GS) was the most abundant enzyme in both oxic and anoxic treatments on the Fe Peak 3 (along the AE600 dimension, Figure 7A and B), yet the GS peak maxima was offset from the iron peak in the size exclusion spectrum at SE 13 in both treatments, implying that perhaps the apo form may have been associated with a larger protein complex. This enzyme is important in nitrogen metabolism that converts glutamate and ammonia into glutamine [53].

*Fe Peak 3b and 3c:* The ribonucleotide reductase subunits NrdAB were the next most abundant iron metalloproteins in the anoxic AE500 and AE 600 dimensions (Figure 7C, H), and the AE 500, but the AE 600 dimensions of the oxic treatment (Figure 7C, G). The ribonucleotide reductases are a key enzyme in DNA replication by providing the necessary dNTP substrates. *P. aeruginosa* is unusual in that it contains all three classes of ribonucleotide reductases within their genomes (RNR I-III) [54,55]. Most RNRs are metalloenzymes, with RNR-I (NrdAB) subclasses having Fe-Fe, Fe-Mn, Mn-Mn, or no metal active sites, RNR-II (NrdJab) requiring B_12_, and RNR-III (NrdDE) requiring two Fe atoms [55–57]. Notably, mutants in the anaerobic RNRs (Class II and III) having deficient biofilm formation^15^ and impaired anaerobic growth and virulence^16^. The factors necessitating this triple redundancy in *Pseudomonas* are thought to be related to hypoxia and are not well understood.

The observation that NrdB shared a maxima with iron in both oxic and anoxic treatments of AE 500 implied this ribonucleotide reductase isoform was a major contributor to the Fe signal at this site, which is known to have 3 Fe atoms per NrdB polypeptide [58]. Interestingly, the distribution of the NrdA subunit overlapped but with a much wider distribution (SE 12-30 in oxic, Figure 7G, and SE12-35 in anoxic, Figure 7H) and had different maxima than NrdB, implying the NrdAB complex may not have been stoichiometrically assembled throughout the dimension. The alignment of NrdB with the Fe Peak in the AE 500 dimension was consistent that the divot in anoxic AE600 SE14 being an artifact of sample processing. The three most abundant iron proteins in this dimension, NrdA, NrdB, and glutamine synthetase, all show adjacent points that imply higher protein abundance if extrapolated between them (e.g. between SE13 and SE15). When linearly extrapolated these three proteins would be the major proteins with maxima close to or potentially on the Fe Peak 3 site (SE 14), implying their contribution to the iron demand here.

The increased contribution of ribonucleotide reductase to Fe Peak 3 was also consistent with the global proteome results (Figures 1 and 8). In our global experiment, the NrdAB is the dominant ribonucleotide reductase under oxic conditions, as expected from prior studies [29]. Under aerobic conditions Class I NrdAB proteins were most abundant, with less of the class II NrdJab, and no observed class III NrdD. Under anaerobic conditions all three classes of RNR’s were present and in higher abundance than in oxic treatments. Similar to CobN, these results also conflict with the model proposed by Crespo et al. (2018), where a cascade of expression from NrdA to NrdJ to NrdG was expected to coincide with decreasing B_12_ availability. Together these results demonstrate a greater role for RNR under anaerobic conditions in *P. aeruginosa* and suggests this enzyme or its cofactors (Fe or B_12_), could be potential antimicrobial targets.

**Figure 8.**
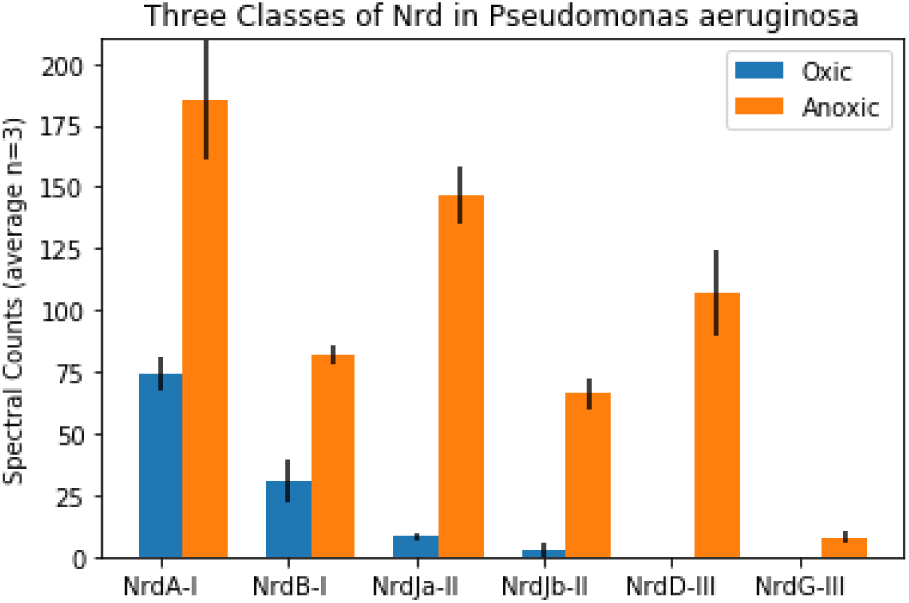
Abundance of proteins associated with the ribonucleotide reductase enzyme complexes within oxic and anoxic treatments from the global proteome data. NrdA/B, and NrdD/G require iron, while NrdJa/Jb requires vitamin B_12_. Bars are the average and error bars represent standard deviation of the extraction triplicates in normalized spectral count units.

The major presence of RNR within the iron metalloproteome and the observation of the triple redundancy of RNR classes present in the proteome under a variety of redox conditions can be attributed to *P. aeruginosa*’s need to maintain high growth rates and corresponding DNA synthesis rates under a variety of oxygen and micronutritional conditions.

*Fe Peak 3d:* A third ferritin annotated as “probably bacterioferritin” (PA4880) was present in the anoxic treatment adjacent to this location with a maximum at SE 18 in the AE 600 dimension (Figure 7D). The ferritins are discussed in greater detail below.

*Summary of Fe Peak 3:* The iron within this peak appeared to be comprised from nitrogen metabolism and DNA synthesis metalloenzymes, as well as co-located iron storage bacterioferritin.

#### Fe Peak 4

Although a relatively minor peak compared to Fe Peaks 1 and 2, the fourth largest peak within the *P. aeruginosa* metalloproteome contained key components of the denitrification apparatus. This peak was found in the AE 200 dimension, at SE-20 and SE-22 in oxic and anoxic treatments, respectively (Table 1, Figures 2 and 3). Similar to Peak 3, the peak was much more pronounced in the anoxic treatment compared to the oxic peak with 7.0 times more Fe in the anoxic at their respective maximum locations (Figures 9G, H). This was large Fe difference occurred despite the anoxic treatment having only 9.2% more total protein (Table S1). Yet Fe Peak 4 was still the smallest of the four iron peaks, with Fe Peak 1 being 5.5 larger in the anoxic treatment. Fe Peak 4 in the oxic treatment was particularly minor, approaching background sounding baseline levels, consistent with minimal need for denitrification apparatus under oxic conditions. Notably, the Fe Peak 4 is quite broad and the known iron metalloproteins within it range from SE 18 to SE 32 (Figures 9B, D, F, and H). The denitrification system is known to be a membrane localized protein complex. Despite the high abundance of some of these proteins, the iron peak was smaller than the other three Fe peaks as mentioned above. This may be a result of underrecovery of the denitrification membrane proteins in the native, non-detergent, system. The elution of Fe Peak 4 in an earlier AE fraction than the other three Fe peaks implies a less negative charge of the proteins found here. Consistent with this, peripheral membrane associated proteins, such as the denitrification and cytochromes enzymes discussed in this section, contain charged surfaces that enables their association with the negative potential of the plasma membrane. [59,60]

**Figure 9.**
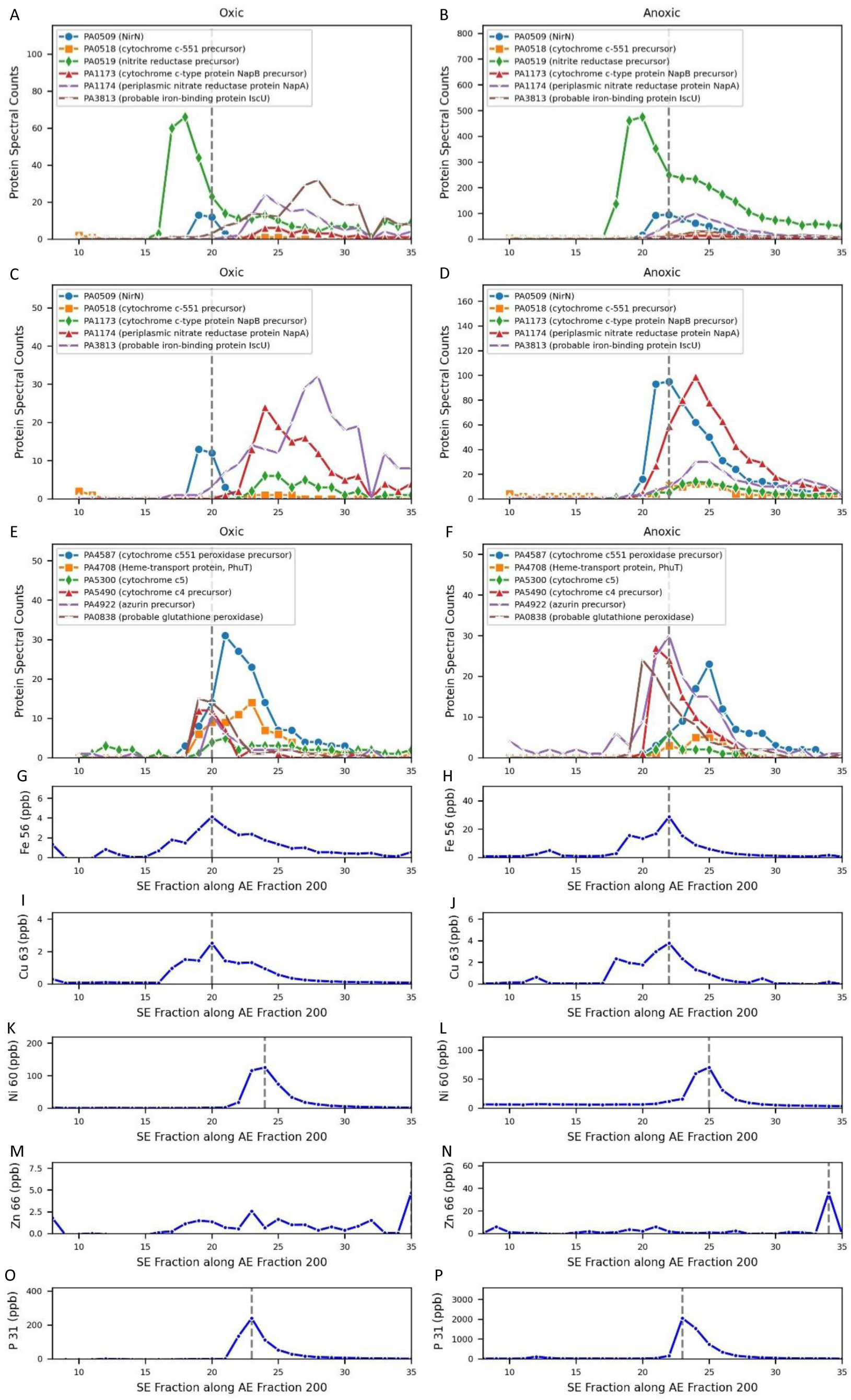
Metalloproteins associated with Fe Peak 4 20 oxic and at anoxic treatments (panels A-F), dashed line is the location of the iron maxima under each condition (AE 200 SE and AE 200 SE 22). Element traces along the AE 200 dimension for Fe, Cu, Ni, Zn, and P traces along the AE 200 dimension (panels G-P), dashed line is the maximum for each element. Left and right panels are oxic and anoxic treatments, respectively. Panels C and D replicate A and B, but without PA0519 to allow examination of lower abundance proteins. Proteins shown are known metalloproteins detected at the iron maxima positions, see Table S2 for list of proteins.

*Proteins in the Anoxic Treatment*: A total of 148 proteins were detected at the anoxic maximum of Fe Peak 4, of those 18 proteins shared a maximum at that location (AE 200 SE-22, Table S2). Of those 18, NirN (PA0509) was the only known iron metalloprotein present and 11 of those were annotated as hypothetical proteins (Figures 9B, D, F). Among the remaining proteins detected at SE-22 (not just having a maxima), 9 were known iron proteins, including components of the denitrification system (Figures 9B, D, F).

*Proteins in the Oxic Treatment*: 162 proteins were detected at the oxic maximum of Fe Peak 4, with 27 sharing a maximum at that location (AE 200 SE-20, Table S2). Of those 27 proteins with a shared maxima none were known iron proteins and 14 were annotated as hypothetical proteins. Among the proteins detected at SE-20 (Figure 9A, C, E), four were known iron proteins, all of which were also on the list of iron proteins detected in the anoxic peak (Table S2), including nitrite reductase precursor (PA0519), cytochrome c4 precursor (PA5490), probable iron-binding protein IscU (PA3813), and cytochrome c551 peroxidase precursor (PA4587).

*Fe Peak 4a-c:* Under anoxic and added nitrate media conditions *Pseudomonas* utilizes denitrification-based respiration. A number of iron proteins are enlisted including NirS (4a: PA0519 nitrite reductase precursor), NapA (4b: periplasmic nitrate reductase protein NapA PA1174), and NirN (4c: PA0509). NirN is a NirS homologue, and forms a 8 blade β-propeller with a cytochrome c domain [61].

Nitrite reductase (NirS, PA0519) was highly abundant in the anoxic treatment. With reaching 475 spectral counts at its maximum, it was the 2^nd^ most protein in the metalloproteomic dataset. While its peak is somewhat offset from the Fe/Cu peak by two SE fractions it still was at 250 spectral counts on the Fe maximum (Figures 9B, J, I).

*Azurin Cu protein:* The copper containing azurin enzyme (PA4922) is an electron donor to nitrite reductase (NirS) [62], and the residues involved in their protein-protein interaction have been identified characterized [63]. Here we observed the Cu peak in the AE 200 dimension to co-elute with Fe Peak 4 in both oxic and anoxic treatments (Figures 9I, J). This protein was also one of the most abundant proteins in the anoxic metalloproteome. While it was present with 30 spectral counts at the maximum in AE 200, it was highly abundant in AE 100 reaching 449 spectral counts at 100-22, where few other proteins elute.

*Fe Peak 4d*: IscU is an Fe-S cluster assembly scaffold, and part of the ISC (Iron Sulfur Cluster) biosynthetic system. [64] Its iron signal may be due to the presence of Fe-S within the protein scaffold. Unlike *E. coli* which has the SUF and ISC Fe-S biosynthesis systems, *P. aeruginosa* only has the ISC system and is essential for viable *Pseudomonas*, has been proposed as a target for antibiotic treatment, although the human mitochondria use a homolog of the system that could be sensitive to treatments.

*Fe Peak 4e:* Corresponding to the occurrence of five cytochromes in this area (see below), a heme-transport protein, PhuT, (PA4708) also co-eluted, consistent with the need to assemble these prosthetic groups into the cytochromes and NirS. PhuS in *P. aeruginosa* has been shown to contribute heme to heme oxygenase and to in sensing and maintaining iron homeostasis. [65] PhuT may have a role in contributing to biosynthesis of denitrification cytochromes and enzymes.

*Fe Peak 4f-j:* Five cytochrome proteins were observed in this range, including cytochrome c4 precursor, cytochrome c-551 precursor, cytochrome c551 peroxidase precursor, cytochrome c5, and cytochrome c-type protein NapB precursor (PA5490, PA0518, PA4587, PA5300, PA1173). Cytochrome proteins help to conduct electrons, they contain the prosthetic group heme, which coordinates an Fe atom at its center. The co-elution of these five cytochromes at the Fe Peak 4 was consistent with their involvement in the denitrification system, and a cytochrome from *Thermus thermophilus* has been proposed to be part of a super complex with nitrate reductase [66].

*Fe Peak 4 - Non-Fe metalloproteins:* In addition to azurin two additional non-iron metalloproteins were detected at Fe Peak 4, nitrous-oxide reductase precursor (PA3392, Cu, anoxic only), probable glutathione peroxidase (PA0838, Se, oxic and anoxic).

*Summary of Fe Peak 4:* This peak was dominated by denitrification metabolism metalloenzymes and associated Fe-S cluster and heme biosynthesis systems and cytochrome electron transport capabilities.

#### Iron Storage Proteins – Bacterioferritin and Ferritin

Given the importance of iron for the variety of metalloenzymes and biochemical functions described above, the ability to store iron when excess is available and to prevent intracellular toxicity is important for bacterial cells. Ferritins are proteins whose 12 or 24 subunits combine to form a protein sphere that stores as many as ∼4000 iron atoms per complex as iron oxide. Bacterioferritins are bacterial versions of ferritin that also differ Eukaryotic ferritins in containing heme between subunits [67]. In a prior application of metalloproteomics on the bacterium *Pseudoalteromonas*, iron storage within two copies of bacterioferritin represented the dominant reservoir of iron within that microbe, indicating a substantial storage capability [21]. Within *P. aeruginosa’s* genome (PAO1) there are three annotated ferritin genes, a bacterioferritin (BfrB, PA3531), a “bacterial ferritin” (FtnA, PA4235), and a probable bacterioferritin (PA4880). There is also a bacterioferritin comigratory protein in the PAO1 genome (PA1008).

Prior structural analysis revealed that *P. aeruginosa* BfrB coordinates heme iron between subunits, while FtnA does not [68], and that a knockout BfrB *P. aeruginosa* strain did not accumulate iron within any other ferritin molecule, leading to their conclusion that BfrB (PA3531) was the main iron storage molecule [69]. Moreover, the *P. aeruginosa* FtnA (PA4235) protein was described as a ferritin distinct from bacterioferritin [70] that assembles as a heterooligomer 24mer from both BfrB and FtnA subunits. A decreasing ratio of BfrB to FtnA was observed at low oxygen, with FtnA becoming the dominant subunit under low oxygen conditions reaching ∼30% to 70% of the subunits [68].

While these first two ferritins (PA3531 and PA4235) have been previously studied, less work has been done on the 2^nd^ bacterioferritin (and third ferritin overall), PA4880. Recent structural studies by Rajapaksha et al [71] showed that PA4880 adopts a 12-mer structure similar to Dps proteins, but lacks the iron coordinating ligands of Dps. Rajapaksha et al. named this gene DpsL (for Dps-like) and suggested that this protein may contribute to avoiding iron toxicity and as well as “innate immune mechanisms consisting of restriction endonucleases and cognate methyl transferases.”

The current metalloproteome study provides a whole organism assessment of the iron storage capabilities in *P. aeruginosa*. Since both the global and metalloproteomic techniques used here measure tryptic peptides, we verified that these three ferritins do not share any tryptic peptides that could cause misidentification from their peptidic constituents (see Figure S4-S6 for sequence alignments). In our metalloproteomes, PA3531 (BfrB) and PA4235 (FtnA) both co-elute with in Fe Peak 1 (Figures 4 and 10) and PA4880 (DpsL) elutes in Fe Peak 3 (Figures 7 and 10). The bacterioferritin-associated ferredoxin Bfd (PA3530) was not identified in either the anoxic or oxic metalloproteome, consistent with observations of Bfd upregulation under low iron conditions and its proposed role in mobilizing iron from bacterioferritin [72].

**Figure 10.**
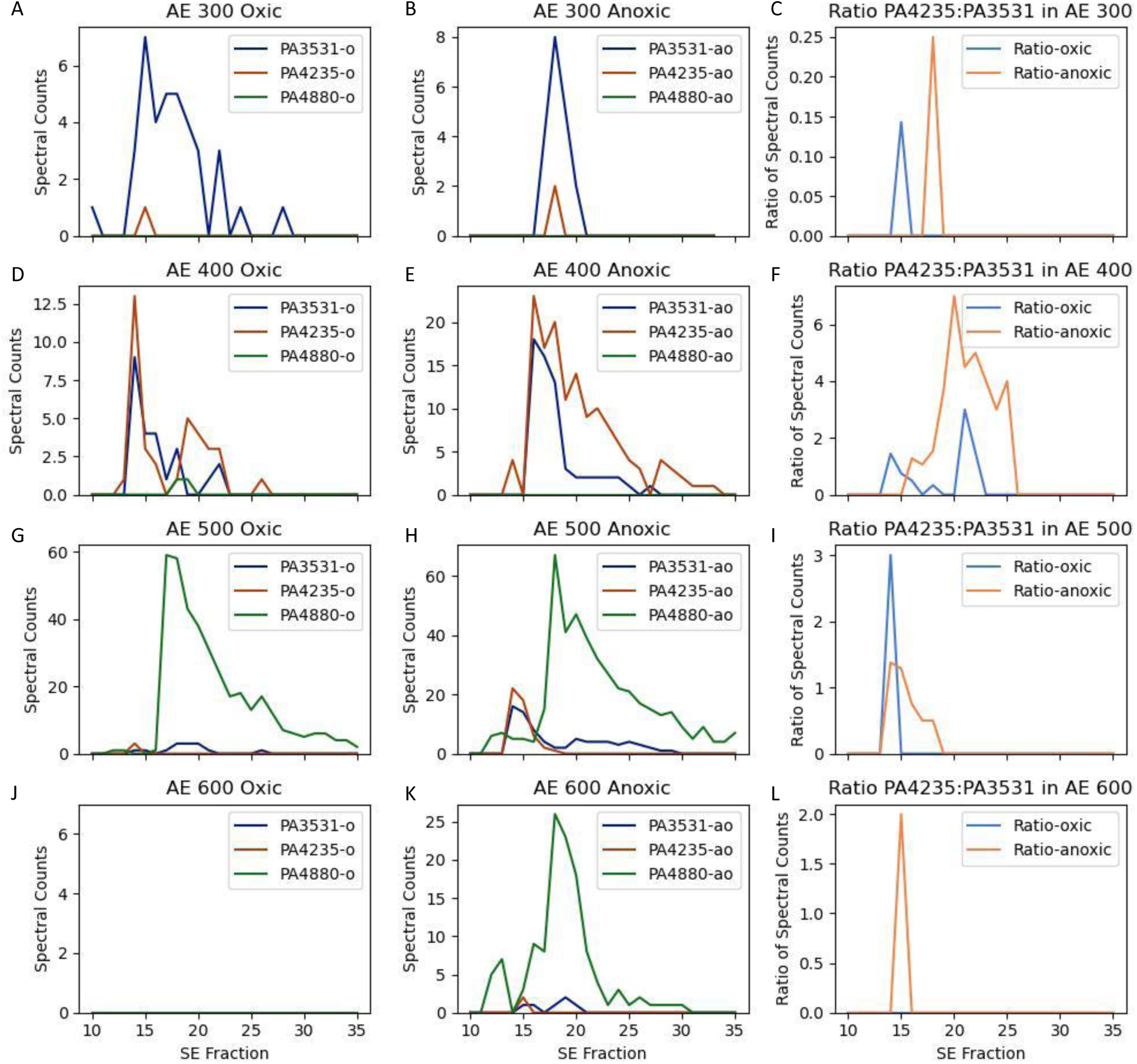
Abundance (oxic: A, D, G, J, anoxic: B, E, H, K; 300, 400, 500, 600) and ratios (C, F, I, L; oxic and anoxic) of ferritin iron storage in *P. aeruginosa*. Distributions of bacterioferritin (PA3531), ferritin (PA4235), and probable bacterioferritin/Dps-like protein (DpsL; PA4880) shown in the 300-600 anion exchange (AE) fractions of the first dimension and 10-35 size exclusion (SE) fractions of the second dimension. PA3531, PA4235, and PA4880 each had the highest spectral counts in AE 300, 400, and 500-600, respectively. Consistent with Yao et al., 2022, the ratio of BfrB:Ftn (PA4235:PA3531; left panels) was highest in the AE 400 anoxic treatment with ratios of 1.1 and 7 between fractions 17 and 25, where much of both ferritins eluted. The PA3531 and PA4235 maxima were slightly offset between treatments (panels D at SE14 in oxic and SE16 in anoxic), likely due to slightly different chromatographies between experiments. No ferritins were observed in the AE 600 oxic fraction.

The hypothesis that the contribution of ferritin (PA4235) increases under anoxic conditions by Yao et al. [68] was supported by our percentage contribution results: PA4235 was the only one of the three proteins to increase in percent contribution across the treatments from 8.2% to 22.2% between oxic and anoxic experiments (Table S1). There also appeared to be support for the accompanying hypothesis that BfrB and FtnA combine as heterogenous subunits to form a single ferritin 24mer protein complex [68]. We observed the ratio of BfrB:Ftn (PA4235:PA3531) to be highest in the AE 400 anoxic treatment with ratios of 1.1 and 7 between fractions 17 and 25 where many of these ferritins eluted (Figure 10F).

Notably the contribution to the native proteome in terms of total spectral counts showed PA4880 to be the most abundant iron storage protein, contrasting with the prior Eshelman et al. (2017) report of PA3531 as the dominant *P. aeruginosa* iron storage molecule as mentioned above. PA4880 had 76.5% of all of the ferritin molecule spectral counts in the oxic treatment and 60.9% in the anoxic treatment (Table S1). PA4880 was the dominant ferritin in AE 500 and AE 600 and coincided with the iron distributions, at SE 17 (oxic) and 18 (anoxic) for AE 500 (Figure 10G and 10H). Notably, PA4880 was not present in oxic AE 600, while it showed a large peak at SE 26 in anoxic AE 600, implying increased use of this bacterioferritin under anoxic conditions. This comparison makes the reasonable assumption that the ionization efficiency of tryptic peptide constituents across these three similar proteins to be approximately averaged out, and that there are similar number of peptides based on their similar small size. More accurate assessments could be conducted in the future by use of targeted proteomics methods [73].

In terms of the amount of iron stored within each of the three ferritins, the metalloproteomic approach could assess this. Yet the high complexity of the *P. aeruginosa* iron metalloproteome makes this less straightforward compared to the straightforward approach used in our prior study on the simpler marine bacterium *Pseudoalteromonas* [21]. From the metalloproteomic data and media conditions of this study, it appears that the extent that the Fe metallome was stored varied considerably between oxic and anoxic treatments. In oxic treatment the two Fe Peak 1 ferritins BfrB and FtnA (PA3531 and PA4235) co-eluted with the major iron peak (Figures 4 and 10), whereas under anoxic conditions they were offset by one higher SE contributing to the shoulder. The extent of iron stored within these three ferritin proteins does not appear to dominate the iron metallome as it did in *Pseudoalteromonas* [21], being a relatively minor peaks compared to the iron metalloproteins, which are offset from the iron maxima, even with the high stoichiometry when fully loaded.

#### Potential Elution of Iron proteins with Protein Assemblages Related to Cellular Metabolisms

Protein assemblies are recognized as a universal feature of cellular biology and are increasingly of interest in the study of subcellular metabolism. In addition to common protein multimers comprised of subunit components, larger assemblies can also exist that including condensates and supercomplexes. Protein condensates, while lacking a consensus definition, include liquid–liquid phase separation (LLPS), and gel and solid assemblies [74], resulting in “membraneless organelles” and resultant functionality [75], and have been invoked within bacteria [76]. Supercomplexes similarly are larger multi-protein assemblies, such as the respiratory complexes associated with the cytoplasmic membrane in the bacterium *Paracoccus denitrificans* [77]. The observation of four Fe peaks in this study in both oxic and anoxic treatments with multiple co-eluting iron protein contributors (Table 1) suggests the potential for the occurrence of multi-protein assemblages in native extracts.

Methodologies for studying protein assemblies have commonalities with metalloproteomic approaches, such employing native size exclusion chromatographic methods [78]. While the native (no detergent) approach used in this study is intended to allow persistence of metal-protein coordination, it also likely contributes to protein complexes having the potential to remain intact within the 2D chromatographic separations. In a study of protein condensates and assemblies, Victor et al additionally employed crosslinking of proteins prior to size exclusion chromatography to fix protein interactions [79]. This methodology employed in this study used minimal sonication that could have contributed to the persistence of some protein assemblies (1 min total on time, see Metalloproteomics methods above), compared to other metalloproteomic studies that used more extensive sonication (12.5 min total on time) that could have disrupted assemblages [24]. Similarly, the presence of protein supercomplexes, and assembles of multiple enzymes complexes (complexes of complexes) has been observed by native gel and chromatographic methods. For example the respiratory complexes, termed respirasomes, within mitochondria and bacteria contain assemblies of respiratory chain complexes III and IV, each containing one or more enzymes [80,81].

To explore this possibility, we calibrated the size exclusion chromatography using commercial standards for protein oligomers (Figure 11A). The first three standards, IgG, BSA (bovine serum albumin) and myoglobin, displayed a linear separation within increasing fraction number as expected with the smaller proteins taking longer to pass through the column due to their interaction with bead pores (linear regression yields an r^2^ of 0.98 and slope of −9.0, Figure 11B). The larger thyroglobulin complex, at 666,000 Da, deviates from this line co-eluting with the IgG at fraction 12, likely related to complexes at this size being too large to interact and be retained by bead pores. Replication of elution fraction was also verified using triplicate separation of *P. aeruginosa* native extract in the 2D metalloproteome separation. The elution of the copper azurin peak showed good reproducibility of elution fraction (Figure 11C). Together these results provide an estimate of the reproducibility and size separation capabilities of the present approach.

**Figure 11.**
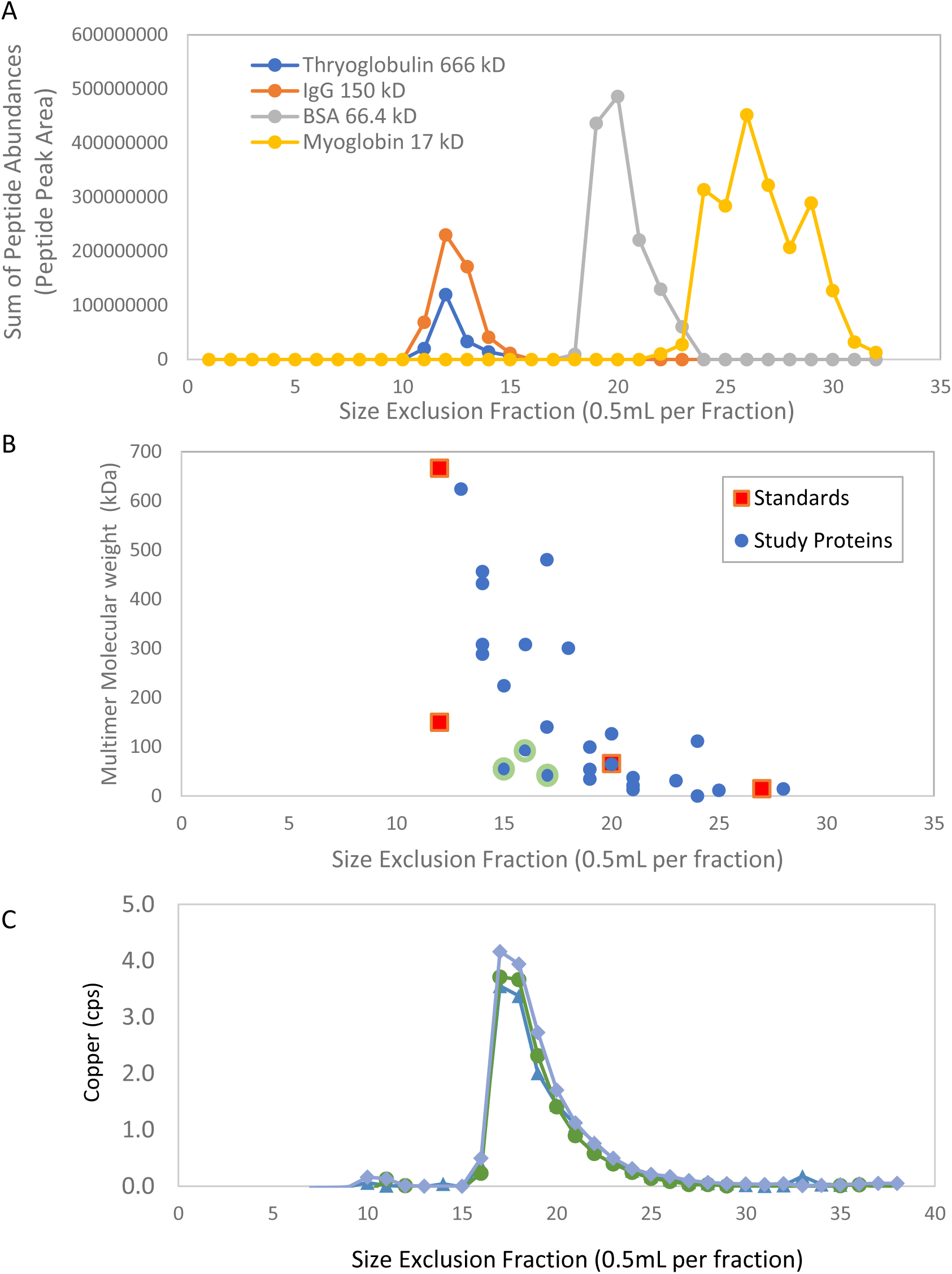
A) Elution of protein size standards on size exclusion column and tryptic digested and analyzed by DIA. The standards throglobulin (666 kD), IgG (150 kD), BSA (66.4 kD), and myoglobin (15 kD) were separated by SE column, then trypsin digested and analyzed by DIA proteomics. B) Comparison of location of peak maxima for protein size standards (from A) and proteins associated with Fe peak 1-4, using multimer molecular weights (Table 2). Three proteins in green circles are hydrazone dehydrogenase (HdhA; PA4022), iron superoxide dismutase (SodB; PA4366), and aconitate hydratase 2 (PA1787) fell below the trend of the three standards: IgG, BSA, and myoglobin (linear regression: r^2^ of 0.98, slope of −9.0). C) Triplicate replicates show reproducible elution of the *P. aeruginosa* metalloprotein azurin, as shown by Cu counts per second (cps). Cu only has one peak in this dimension and hence is useful to verify reproducibility.

The size of the oligomeric forms of iron proteins associated with the four major Fe peaks in this study (Table 2 and references therein for oligomeric form) followed a similar general trend of decreasing size with higher fraction number. However, there was also a wide range of variability in size in each SE fraction (Figure 11B) that could be interpreted as potential association with multi-protein complexes or condensates, particularly in Fe Peak 1-2, whose peaks were eluted at the high molecular weight end of the calibration (SE 14-18; Table 1). For example, three proteins, highlighted in Figure 11B, hydrazone dehydrogenase (HdhA; 55 kDa, PA4022), iron superoxide dismutase (SodB; 21 kDa, PA4366), and aconitate hydratase 2 (Aco2; 94kDa, PA1787), all cytoplasmic enzymes, fell below the calibration standards relationship and most of the other proteins, implying there are protein interactions that result in their earlier SE elution. Pairwise protein-protein interactions of these proteins modeled with Alphafold3 showed potential docking sites between these proteins, with pTM scores of >0.5 but ipTM scores of less than 0.2, suggestive, but not conclusive of interactions that support co-elution observations [82] (structures not shown).

Fe Peak 3 moved from to earlier in the SE elution, from SE 18 to SE 14, between oxic and anoxic treatments (Figure 2; Table 1), implying a larger molecular weight distribution of the much larger Peak 3 iron reservoir present under anoxic conditions. As mentioned above, Fe Peak 3 was associated with the NrdA and NrdB proteins. Notably, NrdA and NrdB have monomer molecular weights of 107 kDa and 47 kDa, while NrdAB forms an active α_2_β_2_ complex and an inactive α_4_β_4_ complex [83], which have weights of 308 kDa and 614 kDa, respectively. Both NrdA and NrdB have wide SE distributions extending from SE 12 to 24 in both the AE 500 and AE 600 dimensions, with NrdA having a two maxima, at SE 13 and SE 23 (AE 500 dimension anoxic), implying the transition between monomer and oligomer forms was captured by the metalloproteomic analysis.

As mentioned above, Fe Peak 4 eluted later in the size exclusion (SE 22) and earlier in the AE fractions (200mM) consistent with less protein-protein interactions, but more positively charged peripheral membrane associated proteins.

Further analysis of co-elution of proteins was conducted by examining the elution of the four iron peaks in relation to the location of the maxima of all identified proteins (Figure 12A) and the sum of all protein abundance at each location (sum of spectral counts at each 2D loci, Figure 12B). The separation of the four iron peaks across four AE fractions (200 mM, 300 mM, 400 mM, and 600 mM), is consistent with distinct protein assemblages in each of those cases. Moreover, two of the four peaks eluted earlier than the location of maximum protein spectral counts in that AE dimension (the 400 mM and 600 mM fractions; Figure 12B), indicating these iron peaks being associated with proteins that were larger than the majority of proteins in that 2^nd^ dimension and consistent multi-protein assemblies present.

**Figure 12.**
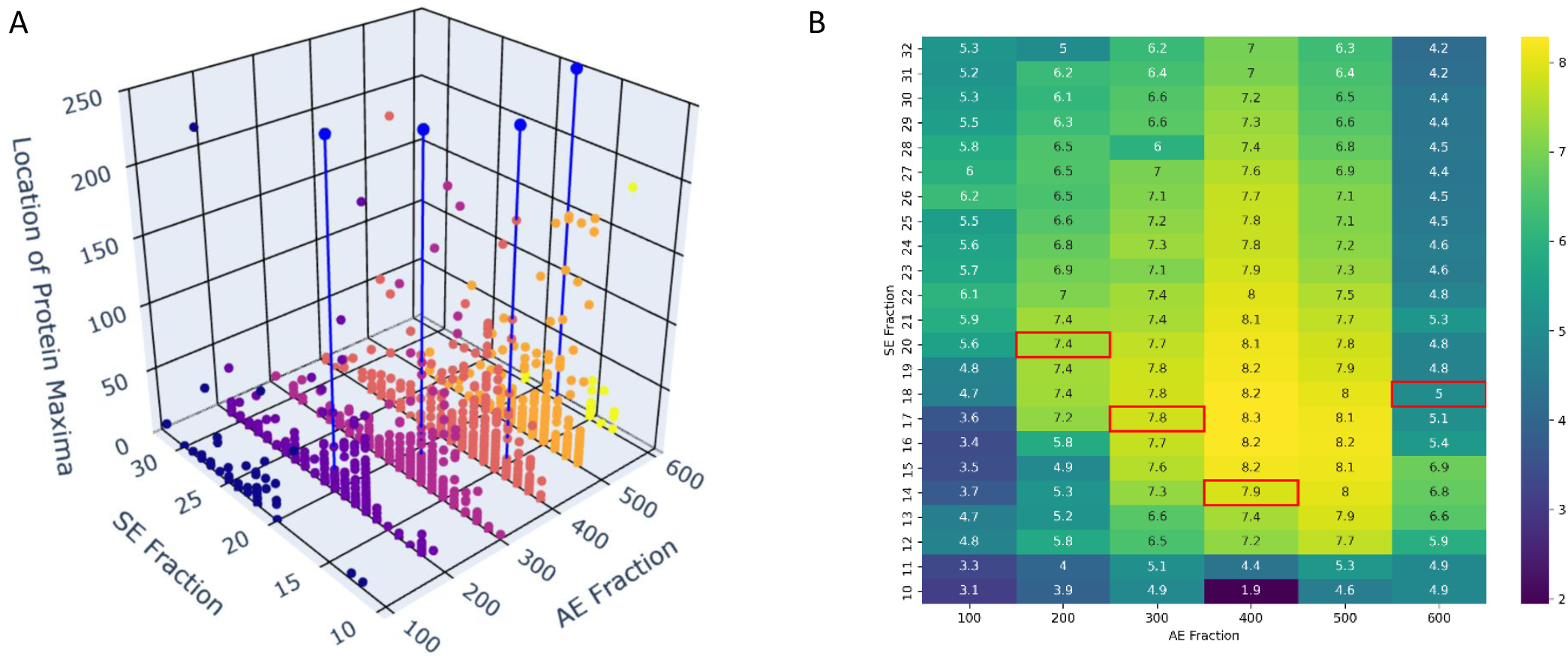
A) Location of the maxima for each protein within the AE-SE 2D chromatographic grid for the oxic treatment, with its associated spectral count value at the maxima (z value) for each protein. Note that each protein is only represented once in the plot. Location of iron peaks shown by blue circles and vertical lines from Table 1. B) Heatmap of log sum of spectral counts of all proteins at each AE and SE fraction location for the oxic treatment. Red boxes indicate the locations of iron peaks from Table 1.

There are interesting implications for potential multi-protein assemblies being present related to iron use within the cytosol of *P. aeruginosa*. In particular, the co-location of proteins with related functions and co-located with dedicated iron storage systems suggests a level of cellular organization at the protein-protein interaction level that may have evolved to be enable some prioritization of metal distribution and trafficking. From a systems biology perspective, the observation of co-eluting and functionally related metalloproteins (Table 2) implies a potential cellular organization combining metabolic functions with iron supply and biosynthesis of Fe-S cluster prosthetic groups within *Pseudomonas*. As bacteria tend to have only a few organelle structures (such as photosynthetic thylakoids, and protein-bound carboxysomes and vacuoles) compared to Eukaryotes [80], the strategies that allow targeting of enzymes and transporting of metals to organelles are less available to them. Yet, the challenges associated with directing metal resources towards specific reactions must be considerable, particularly in rapidly growing and metabolically flexible organisms such as *Pseudomonas* that has competing iron uses. Hence the co-locating of multiple metabolically-related iron enzymes and ferritins storage molecules within “membraneless organelles” of protein condensates would be advantageous, contributing to both the metalation and the prioritization of iron distribution to those metabolisms. The four protein assemblies observed in this study were not necessarily specific to iron metalloproteins: other metalloenzymes and proteins also co-eluted, such as copper enzymes in the Fe Peak 4 region and other non-metalloproteins, hence their subcellular organization may be related to both shared metabolic function and broader metal use.

It is likely that the many protein-protein interactions needed to create these protein assemblies and condensates not only involve a combination of protein binding as briefly explored here, but also the crosstalk of post-translational modifications (PTMs) between proteins. In this latter case, PTMs between proteins may contribute to the associations and functionality. For example, the eukaryotic aconitase enzyme has been shown to undergo reversible oxidation of its iron sulfur cluster and cysteine residues, as well as phosphorylation and transamidation PTMs, all of which contribute to cellular functions [84], and would create a variety of isoforms that elute and associate distinctly. While *in vitro* studies are more common in regulatory PTMs studies [85], *in vivo* datasets such as this native metalloproteomic approach offer an avenue to discover complex biological processes in use and their interaction with other cellular systems. Adding PTM analysis and interpretation of potential protein-protein interactions adds challenges to this already complex metalloproteomic analysis and presents a perhaps daunting level of biological complexity, yet it also presents great opportunities for discovery as this complexity is important to *in vivo* biology. Synergies are possible, such as further use of structural biology to help validate potential PTM sites [81] and protein-protein interactions. Future studies could take advantage of the large volume of raw spectra and their distributions within native separations space, such as from this study and other native datasets, to elucidate PTMs dynamics associated with protein-protein interactions and condensates [77,86].

Future studies could further explore these potential protein assemblies and interrogate the regulation and structural biology behind them. While originally conceived as a methodology capable of investigating metalloproteins, this native 2D approach also serves to contribute empirical information about protein-protein interactions and assembly formation that pertain to cellular metal homeostasis. This study characterized the iron metalloproteome of the more complex prokaryotic microorganisms, attributing enhanced iron use under anaerobic denitrifying metabolism to its specific metalloprotein constituents.

## Data Availability

The processed metalloproteomic metal and protein data, and the global proteomic data are available as supplemental datasets and at Zenodo at 10.5281/zenodo.14950404. For the metalloproteomics four CSV data files are available: 2025_0214_pao_metals_oxic.csv, 2025_0214_pao_metals_anoxic.csv, 2025_0214_pao_proteins_oxic.csv, and 2025_0214_pao_proteins_oxic.csv. For the global proteome one file is provided with average and standard deviation of technical triplicates (pao_df.csv). Raw proteomic spectra are available at ProteomeXchange and PRIDE (review access available upon request). The Jupyter notebook code for data analysis and visualization for this project is available at https://github.com/maksaito/Metalloproteomic-Viewer (Version 12). The *Pseudomonas aeruginosa* genome for the isolate 1-54 is available at Zenodo at 10.5281/zenodo.15336754.

## Acknowledgements

We thank Dawn Moran for microbiological isolations and Viktoria Steck for assistance with genome sequencing samples. We thank Kevin Waldron and Steve Giovannoni for helpful discussions. We also thank two anonymous reviewers for constructive feedback. The research was supported by NIH R01GM135709, the Simons Foundation, and the Center for Chemical Currencies on a Microbial Planet (NSF 2019589; contribution #069), and NSF projects 2123055, 2125063, 2048774.

## Notes

https://zenodo.org/records/14950404

https://zenodo.org/records/15336754

### Competing Interest Statement

The authors have declared no competing interest.

### Summary of Updates

Revision incorporating peer review changes. Final section clarified, final two figures replaced.

## References

1. Palleroni NJ. Prokaryote taxonomy of the 20th century and the impact of studies on the genus*Pseudomonas*: a personal view. Microbiology 2003;149:1–7.

2. Palleroni N, Genius I. *Pseudomonas* Migula 1894. Bergey’s Manual of Systematic Bacteriology 1984;1:141–99.

3. Bodey GP, Bolivar R, Fainstein V et al. Infections caused by *Pseudomonas aeruginosa*. Reviews of Infectious Diseases 1983;5:279–313.

4. Tacconelli E, Carrara E, Savoldi A et al. Discovery, research, and development of new antibiotics: the WHO priority list of antibiotic-resistant bacteria and tuberculosis. The Lancet Infectious Diseases 2018;18:318–327.

5. Skaar EP, Raffatellu M. Metals in infectious diseases and nutritional immunity. Metallomics 2015;7:926–8.

6. Kimata N, Nishino T, Suzuki S et al. *Pseudomonas aeruginosa* isolated from marine environments in Tokyo Bay. Microb Ecol 2004;47:41–7.

7. Crone S, Vives-Flórez M, Kvich L et al. The environmental occurrence of *Pseudomonas aeruginosa*. APMIS 2020;128:220–31.

8. Borer B, Zhang IH, Baker AE et al. Porous marine snow differentially benefits chemotactic, motile, and nonmotile bacteria. PNAS Nexus 2023;2:pgac311.

9. Carlson CA, Ingraham JL. Comparison of denitrification by *Pseudomonas stutzeri*, Pseudomonas aeruginosa, and Paracoccus denitrificans. Appl Environ Micro 1983;45:1247–1253.

10. Thomas KL, Lloyd D, Boddy L. Effects of oxygen, pH and nitrate concentration on denitrification by *Pseudomonas* species. FEMS Microbiol Lett 1994;118:181–186.

11. Jesaitis AJ, Franklin MJ, Berglund D et al. Compromised host defense on *Pseudomonas aeruginosa* biofilms: characterization of neutrophil and biofilm interactions. J Immunol 2003;171:4329–4339.

12. Poulsen BE, Yang R, Clatworthy AE et al. Defining the core essential genome of *Pseudomonas aeruginosa*. Proc Natl Acad Sci U S A 2019;116:10072–80.

13. Stover C, Pham X, Erwin A et al. Complete genome sequence of *Pseudomonas aeruginosa* PAO1, an opportunistic pathogen. Nature 2000;406:959–64.

14. Golden MM, Heppe AC, Zaremba CL et al. Metal Chelation as an Antibacterial Strategy for *Pseudomonas aeruginosa* and *Acinetobacter baumannii*. RSC Chem Biol 2024, DOI: 10.1039/D4CB00175C.

15. Soldano A, Yao H, Punchi Hewage AN et al. Small Molecule Inhibitors of the Bacterioferritin (BfrB)– Ferredoxin (Bfd) Complex Kill Biofilm-Embedded *Pseudomonas aeruginosa* Cells. ACS Infectious Diseases 2020;7:123–40.

16. Sánchez-Jiménez A, Marcos-Torres FJ, Llamas MA. Mechanisms of iron homeostasis in *Pseudomonas aeruginosa* and emerging therapeutics directed to disrupt this vital process. Microb Biotechnol 2023;16:1475–91.

17. Schalk IJ, Perraud Q. *Pseudomonas aeruginosa* and its multiple strategies to access iron. Environ Microbiol 2023;25:811–31.

18. Mastropasqua MC, D’Orazio M, Cerasi M et al. Growth of *Pseudomonas aeruginosa* in zinc poor environments is promoted by a nicotianamine-related metallophore. Mol Microbiol 2017;106:543–561.

19. Barnett JP, Scanlan DJ, Blindauer CA. Fractionation and identification of metalloproteins from a marine cyanobacterium. Anal Bioanal Chem 2012;402:3371–7.

20. Cvetkovic A, Menon AL, Thorgersen MP et al. Microbial metalloproteomes are largely uncharacterized. Nature 2010;466:779–82.

21. Mazzotta MG, McIlvin MR, Saito MA. Characterization of the Fe metalloproteome of a ubiquitous marine heterotroph, *Pseudoalteromonas* (BB2-AT2): multiple bacterioferritin copies enable significant Fe storage. Metallomics 2020;12:654–67.

22. Mazzotta MG, McIlvin MR, Moran DM et al. Characterization of the metalloproteome of *Pseudoalteromonas* (BB2-AT2): Biogeochemical underpinnings for zinc, manganese, cobalt, and nickel cycling in a ubiquitous marine heterotroph. Metallomics 2021;13:mfab060.

23. Tottey S, Patterson CJ, Banci L et al. Cyanobacterial metallochaperone inhibits deleterious side reactions of copper. Proc Nat Acad Sci USA 2012;109:95–100.

24. Neville SL, Eijkelkamp BA, Lothian A et al. Cadmium stress dictates central carbon flux and alters membrane composition in *Streptococcus pneumoniae*. Communications Biol 2020;3:694.

25. Saunders JK, McIlvin MR, Dupont CL et al. Microbial functional diversity across biogeochemical provinces in the central Pacific Ocean. Proc Nat Acad Sci USA 2022;119:e2200014119.

26. Cohen NR, McIlvin MR, Moran DM et al. Dinoflagellates alter their carbon and nutrient metabolic strategies across environmental gradients in the central Pacific Ocean. Nature Microbiol 2021;6:173–86.

27. Rodionov DA, Vitreschak AG, Mironov AA et al. Comparative genomics of the vitamin B_12_ metabolism and regulation in prokaryotes. J Bacteriol 2003;278:41148–59.

28. Teitzel GM, Geddie A, De Long SK et al. Survival and growth in the presence of elevated copper: transcriptional profiling of copper-stressed *Pseudomonas aeruginosa*. J Bacteriol 2006;188:7242–56.

29. Crespo A, Blanco-Cabra N, Torrents E. Aerobic Vitamin B_12_ Biosynthesis Is Essential for *Pseudomonas aeruginosa* Class II Ribonucleotide Reductase Activity During Planktonic and Biofilm Growth. Frontiers Microbiol 2018;9:986.

30. Dunwell JM, Purvis A, Khuri S. Cupins: the most functionally diverse protein superfamily? Phytochem 2004;65:7–17.

31. Harmer CJ, Wynn M, Pinto R et al. Homogentisate 1-2-dioxygenase downregulation in the chronic persistence of *Pseudomonas aeruginosa* Australian epidemic strain-1 in the CF lung. PLoS One 2015;10:e0134229.

32. Ascher DB, Spiga O, Sekelska M et al. Homogentisate 1,2-dioxygenase (HGD) gene variants, their analysis and genotype-phenotype correlations in the largest cohort of patients with AKU. Eur J Hum Genet 2019;27:888–902.

33. Hassett DJ, Sutton MD, Schurr MJ et al. *Pseudomonas aeruginosa* hypoxic or anaerobic biofilm infections within cystic fibrosis airways. Trends Microbiol 2009;17:130–8.

34. Taniyama K, Itoh H, Takuwa A et al. Group X aldehyde dehydrogenases of *Pseudomonas aeruginosa* PAO1 degrade hydrazones. J Bacteriol 2012;194:1447–56.

35. Castro L, Tórtora V, Mansilla S et al. Aconitases: Non-redox iron–sulfur proteins sensitive to reactive species. Accounts of Chem Res 2019;52:2609–19.

36. Hassett DJ, Woodruff WA, Wozniak DJ et al. Cloning and characterization of the *Pseudomonas aeruginosa* sodA and sodB genes encoding manganese- and iron-cofactored superoxide dismutase: demonstration of increased manganese superoxide dismutase activity in alginate-producing bacteria. J Bacteriol 1993;175:7658–65.

37. Polack B, Dacheux D, Delic-Attree I et al. Role of manganese superoxide dismutase in a mucoid isolate of *Pseudomonas aeruginosa*: adaptation to oxidative stress. Infect Immun 1996;64:2216–9.

38. Nishida-Tamehiro K, Kimura A, Tsubata T et al. Antioxidative enzyme NAD (P) H quinone oxidoreductase 1 (NQO1) modulates the differentiation of Th17 cells by regulating ROS levels. Plos One 2022;17:e0272090.

39. Yagi T. Bacterial NADH-quinone oxidoreductases. J Bioenergetics Biomembranes 1991;23:211–25.

40. Pey AL, Megarity CF, Timson DJ. NAD(P)H quinone oxidoreductase (NQO1): an enzyme which needs just enough mobility, in just the right places. Bioscience Reports 2019;39:BSR20180459–BSR20180459.

41. Baillon M-LA, van Vliet AH, Ketley JM et al. An iron-regulated alkyl hydroperoxide reductase (AhpC) confers aerotolerance and oxidative stress resistance to the microaerophilic pathogen *Campylobacter jejuni*. J Bacteriol 1999;181:4798–804.

42. Saunders JK, Gaylord DA, Held NA et al. METATRYP v 2.0: Metaproteomic least common ancestor analysis for taxonomic inference using specialized sequence assemblies—standalone software and web servers for marine microorganisms and coronaviruses. J Proteome Res 2020;19:4718–29.

43. Seaver LC, Imlay JA. Alkyl hydroperoxide reductase is the primary scavenger of endogenous hydrogen peroxide in Escherichia coli. J Bacteriol 2001;183:7173–81.

44. da Cruz Nizer WS, Inkovskiy V, Versey Z et al. Oxidative stress response in *Pseudomonas aeruginosa*. Pathogens 2021;10:1187.

45. Hare NJ, Scott NE, Shin EHH et al. Proteomics of the oxidative stress response induced by hydrogen peroxide and paraquat reveals a novel AhpC-like protein in Pseudomonas aeruginosa. Proteomics 2011;11:3056–69.

46. Lim CK, Hassan KA, Tetu SG et al. The effect of iron limitation on the transcriptome and proteome of *Pseudomonas fluorescens* Pf-5. PLoS one 2012;7:e39139.

47. Ma J-F, Ochsner UA, Klotz MG et al. Bacterioferritin A modulates catalase A (KatA) activity and resistance to hydrogen peroxide in *Pseudomonas aeruginosa*. J Bacteriol 1999;181:3730–42.

48. Lee JT, Lee SS, Mondal S et al. Enhancement of the Chaperone Activity of Alkyl Hydroperoxide Reductase C from *Pseudomonas aeruginosa* PAO1 Resulting from a Point-Specific Mutation Confers Heat Tolerance in *Escherichia coli*. Mol Cells 2016;39:594–602.

49. Arai T, Kimata S, Mochizuki D et al. NADH oxidase and alkyl hydroperoxide reductase subunit C (peroxiredoxin) from *Amphibacillus xylanus* form an oligomeric assembly. FEBS Open Bio 2015;5:124–31.

50. Ochsner UA, Hassett DJ, Vasil ML. Genetic and physiological characterization of ohr, encoding a protein involved in organic hydroperoxide resistance in *Pseudomonas aeruginosa*. J Bacteriol 2001;183:773–8.

51. Fuangthong M, Helmann JD. The OhrR repressor senses organic hydroperoxides by reversible formation of a cysteine-sulfenic acid derivative. Proc Nat Acad Sci USA 2002;99:6690–5.

52. Chen L, Xie QW, Nathan C. Alkyl hydroperoxide reductase subunit C (AhpC) protects bacterial and human cells against reactive nitrogen intermediates. Mol Cell 1998;1:795–805.

53. Brown-Grant K, Tata J. The distribution and metabolism of thyroxine and 3: 5: 3′-triiodothyronine in the rabbit. J Physiol 1961;157:157.

54. Borovok I, Gorovitz B, Yanku M et al. Alternative oxygen-dependent and oxygen-independent ribonucleotide reductases in *Streptomyces*: cross-regulation and physiological role in response to oxygen limitation. Mol Microbiol 2004;54:1022–1035.

55. Jordan A, Torrents E, Sala I et al. Ribonucleotide reduction in *Pseudomonas* species: simultaneous presence of active enzymes from different classes. J Bacteriol 1999;181:3974–3980.

56. Blaesi EJ, Palowitch GM, Hu K et al. Metal-free class Ie ribonucleotide reductase from pathogens initiates catalysis with a tyrosine-derived dihydroxyphenylalanine radical. Proc Nat Acad Sci USA 2018;115:10022–10027.

57. Stubbe J, Seyedsayamdost MR. Discovery of a New Class I Ribonucleotide Reductase with an Essential DOPA Radical and NO Metal as an Initiator of Long-Range Radical Transfer. ACS Publications 2018.

58. Torrents E, Westman M, Sahlin M et al. Ribonucleotide reductase modularity: Atypical duplication of the ATP-cone domain in *Pseudomonas aeruginosa*. J Biol Chem 2006;281:25287–96.

59. Kyne C, Jordon K, Filoti DI et al. Protein charge determination and implications for interactions in cell extracts. Protein Science 2017;26:258–67.

60. Ma Y, Poole K, Goyette J et al. Introducing membrane charge and membrane potential to T cell signaling. Front Immunol. 2017. 2017.

61. Klünemann T, Preuß A, Adamczack J et al. Crystal Structure of Dihydro-Heme d1 Dehydrogenase NirN from *Pseudomonas aeruginosa* Reveals Amino Acid Residues Essential for Catalysis. J Mol Biol 2019;431:3246–60.

62. Kakutani T, Watanabe H, Arima K et al. A blue protein as an inactivating factor for nitrite reductase from *Alcaligenes faecalis* strain S-6. J Biochem 1981;89:463–72.

63. Kukimoto M, Nishiyama M, Tanokura M et al. Studies on protein-protein interaction between copper-containing nitrite reductase and pseudoazurin from *Alcaligenes faecalis* S-6. J Biol Chem 1996;271:13680–3.

64. Romsang A, Dubbs JM, Mongkolsuk S. The Iron–sulfur Cluster Biosynthesis Regulator IscR Contributes to Iron Homeostasis and Resistance to Oxidants in *Pseudomonas aeruginosa*. Stress and Environmental Regulation of Gene Expression and Adaptation in Bacteria 2016:1090–102.

65. Kaur AP, Lansky IB, Wilks A. The role of the cytoplasmic heme-binding protein (PhuS) of *Pseudomonas aeruginosa* in intracellular heme trafficking and iron homeostasis. J Biol Chem 2009;284:56–66.

66. Cava F, Zafra O, Berenguer J. A cytochrome c containing nitrate reductase plays a role in electron transport for denitrification in *Thermus thermophilus* without involvement of the bc respiratory complex. Mol Microbiol 2008;70:507–18.

67. Grossman MJ, Hinton SM, Minak-Bernero V et al. Unification of the ferritin family of proteins. Proc Natl Acad Sci U S A 1992;89:2419–23.

68. Yao H, Soldano A, Fontenot L et al. *Pseudomonas aeruginosa* Bacterioferritin Is Assembled from FtnA and BfrB Subunits with the Relative Proportions Dependent on the Environmental Oxygen Availability. Biomolecules 2022;12:366.

69. Eshelman K, Yao H, Punchi Hewage AND et al. Inhibiting the BfrB:Bfd interaction in Pseudomonas aeruginosa causes irreversible iron accumulation in bacterioferritin and iron deficiency in the bacterial cytosol. Metallomics 2017;9:646–59.

70. Yao H, Jepkorir G, Lovell S et al. Two distinct ferritin-like molecules in *Pseudomonas aeruginosa*: the product of the bfrA gene is a bacterial ferritin (FtnA) and not a bacterioferritin (Bfr). Biochemistry 2011;50:5236–48.

71. Rajapaksha N, Yao H, Cook A et al. Pseudomonas aeruginosa gene PA4880 encodes a Dps-like protein with a Dps fold, bacterioferritin-type ferroxidase centers, and endonuclease activity. Front Mol Biosci 2024;11:1390745.

72. Wang Y, Yao H, Cheng Y et al. Characterization of the bacterioferritin/bacterioferritin associated ferredoxin protein–protein interaction in solution and determination of binding energy hot spots. Biochemistry 2015;54:6162–75.

73. Saito MA, McIlvin MR, Moran DM et al. Abundant nitrite-oxidizing metalloenzymes in the mesopelagic zone of the tropical Pacific Ocean. Nature Geo 2020;13:355–62.

74. Wang H-W. Introduction to the Protein Condensates Virtual Special Issue. Biochemistry 2022;61:2441–2.

75. Mohanty P, Kapoor U, Sundaravadivelu Devarajan D et al. Principles governing the phase separation of multidomain proteins. Biochemistry 2022;61:2443–55.

76. Cohan MC, Pappu RV. Making the case for disordered proteins and biomolecular condensates in bacteria. Trends Biochem Sci 2020;45:668–80.

77. Stroh A, Anderka O, Pfeiffer K et al. Assembly of respiratory complexes I, III, and IV into NADH oxidase supercomplex stabilizes complex I in *Paracoccus denitrificans*. J Biol Chem 2004;279:5000–7.

78. Fekete S, Beck A, Veuthey J-L et al. Theory and practice of size exclusion chromatography for the analysis of protein aggregates. J Pharm Biomedical Analysis 2014;101:161–73.

79. Victor RA, Thompson VF, Schwartz JC. Isolating and analyzing protein containing granules from cells. Current Protocols 2021;1:e35.

80. Murat D, Byrne M, Komeili A. Cell biology of prokaryotic organelles. Cold Spring Harbor perspectives in Biology 2010;2:a000422.

81. Vandermarliere E, Martens L. Protein structure as a means to triage proposed PTM sites. Proteomics 2013;13:1028–35.

82. Lin Y, Wallis C, Corry B. AlphaFold can be used to predict the oligomeric states of proteins. bioRxiv 2025:2025–03.

83. Ando N, Brignole EJ, Zimanyi CM et al. Structural interconversions modulate activity of *Escherichia coli* ribonucleotide reductase. Proc Nat Acad Sci USA 2011;108:21046–51.

84. Oleh V. Lushchak FG Marta Piroddi, Lushchak VI. Aconitase post-translational modification as a key in linkage between Krebs cycle, iron homeostasis, redox signaling, and metabolism of reactive oxygen species. Redox Report 2014;19:8–15.

85. Venne AS, Kollipara L, Zahedi RP. The next level of complexity: crosstalk of posttranslational modifications. Proteomics 2014;14:513–24.

86. Vartak R, Porras CA-M, Bai Y. Respiratory supercomplexes: structure, function and assembly. Protein & Cell 2013;4:582–90.

87. Akram M, Dietl A, Mersdorf U et al. A 192-heme electron transfer network in the hydrazine dehydrogenase complex. Sci Advances 2019;5:eaav4310.

88. Sheng Y, Abreu IA, Cabelli DE et al. Superoxide dismutases and superoxide reductases. Chemical Reviews 2014;114:3854–918.

89. Yang C, Huang Z, Zhang X, et al. Structural insights into the NAD (P) H: quinone oxidoreductase from *Phytophthora capsici*. ACS Omega 2022;7:25705–14.

90. Joo HK, Park YW, Jang YY et al. Structural analysis of glutamine synthetase from *Helicobacter pylori*. Scientific Reports 2018;8:11657.

91. Nurizzo D, Silvestrini M-C, Mathieu M et al. N-terminal arm exchange is observed in the 2.15 Å crystal structure of oxidized nitrite reductase from *Pseudomonas aeruginosa*. Structure 1997;5:1157–71.

92. Henriques BJ, Olsen RKJ, Gomes CM et al. Electron transfer flavoprotein and its role in mitochondrial energy metabolism in health and disease. Gene 2021;776:145407.

93. Nader S, Pérard J, Carpentier P et al. New insights into the tetrameric family of the Fur metalloregulators. Biometals 2019;32:501–19.

94. Liu L, Jiang S, Xing M et al. Structural analysis of an L-cysteine desulfurase from an Ssp DNA phosphorothioation system. mBio 11: e00488–20. 2020.

95. Jacques JG, Fourmond V, Arnoux P et al. Reductive activation in periplasmic nitrate reductase involves chemical modifications of the Mo-cofactor beyond the first coordination sphere of the metal ion. Biochimica et Biophysica Acta (BBA)-Bioenergetics 2014;1837:277–86.

96. Coelho C, González PJ, Moura JG et al. The crystal structure of Cupriavidus necator nitrate reductase in oxidized and partially reduced states. J Mol Biol 2011;408:932–48.

97. Adamczack J, Hoffmann M, Papke U et al. NirN protein from *Pseudomonas aeruginosa* is a novel electron-bifurcating dehydrogenase catalyzing the last step of heme d1 biosynthesis. J Biol Chem 2014;289:30753–62.

98. Kim JH, Tonelli M, Kim T et al. Three-dimensional structure and determinants of stability of the iron– sulfur cluster scaffold protein IscU from *Escherichia coli*. Biochemistry 2012;51:5557–63.

99. Ho WW, Li H, Eakanunkul S et al. Holo-and apo-bound structures of bacterial periplasmic heme-binding proteins. J Biol Chem 2007;282:35796–802.

100. Kadziola A, Larsen S. Crystal structure of the dihaem cytochrome c4 from *Pseudomonas stutzeri* determined at 2.2 Å resolution. Structure 1997;5:203–16.

101. Nordling M, Young S, Karlsson BG et al. The structural gene for cytochrome c551 from *Pseudomonas aeruginosa*: the nucleotide sequence shows a location downstream of the nitrite reductase gene. FEBS letters 1990;259:230–2.

102. Matsuura Y, Takano T, Dickerson RE. Structure of cytochrome c551 from *Pseudomonas aeruginosa* refined at 1.6 Å resolution and comparison of the two redox forms. J Mol Biol 1982;156:389–409.

103. Brown K, Djinovic-Carugo K, Haltia T et al. Revisiting the catalytic CuZ cluster of nitrous oxide (N2O) reductase: evidence of a bridging inorganic sulfur. J Biol Chem 2000;275:41133–6.

104. Nar H, Messerschmidt A, Huber R et al. Crystal structure of *Pseudomonas aeruginosa* apo-azurin at 1.85 Å resolution. FEBS letters 1992;306:119–24.

