## Supplemental Materials for "The Iron Metalloproteome of *Pseudomonas aeruginosa* Under Oxic and Anoxic Conditions"

***For "Detection of Iron Protein Supercomplexes in Pseudomonas aeruginosa by Native Metalloproteomics"***

**Version 5/15/2025**

**Table S1.** $1^{st}$ and $2^{nd}$ dimension metalloproteome samples. Note that analyses have different extents of coverage of the $2^{nd}$ fractions. Matrices were created with no data (not a number, NAN) fields and visualizations adjusted depending on if the extra fractions were to be included or not.

| Anion Exchange Fraction ID Number | NaCl mM (labeled as) | Total Protein in AE Oxic (µg/µL) | Total Protein in AE Anoxic (µg/µL) | % Difference between Anoxic – Oxic Total Protein | $2^{nd}$ Dimension Fractions Metals Oxic | $2^{nd}$ Dimension Fractions Metals Anoxic | $2^{nd}$ Dimension Fractions Proteins Oxic | $2^{nd}$ Dimension Fractions Proteins Anoxic |
|---|---|---|---|---|---|---|---|---|
| 1 | 0 | 0.045 | 0.054 | 20.7 | ND | ND | ND | ND |
| 2 | 0-100 | 0.019 | 0.025 | 37.5 | ND | ND | ND | ND |
| 3 | 100-200 (100) | 0.131 | 0.205 | 56.5 | 7-38 | 7-38 | 10-35 | 10-35 |
| 4 | 200-300 (200) | 0.476 | 0.520 | 9.2 | 7-38 | 7-38 | 10-35 | 10-35 |
| 5 | 300-400 (300) | 0.604 | 0.761 | 26.1 | 7-38 | 7-38 | 10-35 | 10-35 |
| 6 | 400-500 (400) | 1.125 | 1.239 | 10.2 | 7-38 | 7-38 | 10-35 | 10-35 |
| 7 | 500-600 (500) | 0.838 | 1.102 | 31.5 | 7-38 | 7-38 | 10-35 | 10-35 |
| 8 | 600-800 (600) | 0.227 | 0.279 | 23.1 | 7-38 | 7-38 | 10-35 | 10-35 |
| 9 | 800-1000 (800) | 0.093 | 0.130 | 40.8 | 7-38 | ND | ND | ND |
| 10 | 1000-1000 (1000) | 0.042 | 0.063 | 49.1 | 7-38 | ND | ND | ND |
| Number of samples | -- | | | | 256 | 192 | 156 | 156 |

**Table S2**. Proteins present in *P. aeruginosa* metalloproteome Fe Peak 4 under oxic and anoxic conditions

| PA ID | Annotation | Protein Present | Comment |
|---|---|---|---|
| *Putative Iron Proteins* | | | |
| PA0509 | NirN | anoxic | Maxima on anoxic peak |
| PA0519 | nitrite reductase precursor | anoxic/oxic | -- |
| PA1174 | periplasmic nitrate reductase protein NapA | anoxic | -- |
| PA5490 | cytochrome c4 precursor | anoxic/oxic | -- |
| PA0518 | cytochrome c-551 precursor | anoxic | -- |
| PA3813 | probable iron-binding protein IscU | anoxic/oxic | -- |
| PA4587 | cytochrome c551 peroxidase precursor | anoxic/oxic | -- |
| PA5300 | cytochrome c5 | anoxic | -- |
| PA1173 | cytochrome c-type protein NapB precursor | anoxic | -- |
| PA4708 | Heme-transport protein, PhuT | anoxic | -- |
| *Other Metalloproteins* | | | |
| PA3392 | nitrous-oxide reductase precursor | anoxic | -- |
| PA4922 | azurin precursor | anoxic/oxic | -- |
| PA0838 | probable glutathione peroxidase | anoxic/oxic | -- |
| *Other Proteins with maxima on AE 200 SE 20 (oxic) or AE 200 SE 22 (anoxic)* | | | |
| PA3653 | ribosome recycling factor | oxic | Maximum on oxic peak |
| PA2532 | thiol peroxidase | oxic | Maximum on oxic peak |
| PA4611 | hypothetical protein | oxic | Maximum on oxic peak |
| PA2980 | conserved hypothetical protein | oxic | Maximum on oxic peak |
| PA3202 | conserved hypothetical protein | oxic | Maximum on oxic peak |
| PA1810 | NppA2 | oxic | Maximum on oxic peak |
| PA0591 | conserved hypothetical protein | oxic | Maximum on oxic peak |
| PA2134 | hypothetical protein | oxic | Maximum on oxic peak |
| PA4529 | dephosphocoenzyme A kinase | oxic | Maximum on oxic peak |
| PA3383 | binding protein component of ABC phosphonate t... | oxic | Maximum on oxic peak |
| PA3270 | hypothetical protein | oxic | Maximum on oxic peak |
| PA1250 | alkaline proteinase inhibitor AprI | oxic | Maximum on oxic peak |
| PA1749 | hypothetical protein | oxic | Maximum on oxic peak |
| PA0350 | dihydrofolate reductase | oxic | Maximum on oxic peak |
| PA0025 | shikimate dehydrogenase | oxic | Maximum on oxic peak |
| PA0415 | probable chemotaxis protein | oxic | Maximum on oxic peak |
| PA4913 | probable binding protein component of ABC tran... | oxic | Maximum on oxic peak |
| PA0741 | conserved hypothetical protein | oxic | Maximum on oxic peak |
| PA4616 | probable c4-dicarboxylate-binding protein | oxic | Maximum on oxic peak |
| PA2412 | conserved hypothetical protein | oxic | Maximum on oxic peak |
| PA0339 | hypothetical protein | oxic | Maximum on oxic peak |
| PA1890 | probable glutathione S-transferase | oxic | Maximum on oxic peak |
| PA4543 | conserved hypothetical protein | oxic | Maximum on oxic peak |
| PA3801 | conserved hypothetical protein | oxic | Maximum on oxic peak |

| | | | |
|---|---|---|---|
| PA3576 | hypothetical protein | oxic | Maximum on oxic peak |
| PA4164 | hypothetical protein | oxic | Maximum on oxic peak |
| PA0517 | probable c-type cytochrome precurso | oxic | Maximum on oxic peak |
| PA3655 | elongation factor Ts | anoxic | Maximum on anoxic peak |
| PA2442 | glycine cleavage system protein T2 | anoxic | Maximum on anoxic peak |
| PA4738 | conserved hypothetical protein | anoxic | Maximum on anoxic peak |
| PA4611 | hypothetical protein | anoxic | Maximum on anoxic peak |
| PA3785 | conserved hypothetical protein | anoxic | Maximum on anoxic peak |
| PA3182 | 6-phosphogluconolactonase | anoxic | Maximum on anoxic peak |
| PA4463 | conserved hypothetical protein | anoxic | Maximum on anoxic peak |
| PA1123 | hypothetical protein | anoxic | Maximum on anoxic peak |
| PA0741 | conserved hypothetical protein | anoxic | Maximum on anoxic peak |
| PA0950 | probable arsenate reductase | anoxic | Maximum on anoxic peak |
| PA5381 | hypothetical protein | anoxic | Maximum on anoxic peak |
| PA0380 | conserved hypothetical protein | anoxic | Maximum on anoxic peak |
| PA4866 | conserved hypothetical protein | anoxic | Maximum on anoxic peak |
| PA1946 | binding protein component precursor of ABC rib… | anoxic | Maximum on anoxic peak |
| PA1040 | hypothetical protein | anoxic | Maximum on anoxic peak |
| PA2792 | hypothetical protein | anoxic | Maximum on anoxic peak |
| PA3976 | thiamin-phosphate pyrophosphorylase | anoxic | Maximum on anoxic peak |
| PA0283 | sulfate-binding protein precursor | anoxic | Maximum on anoxic peak |

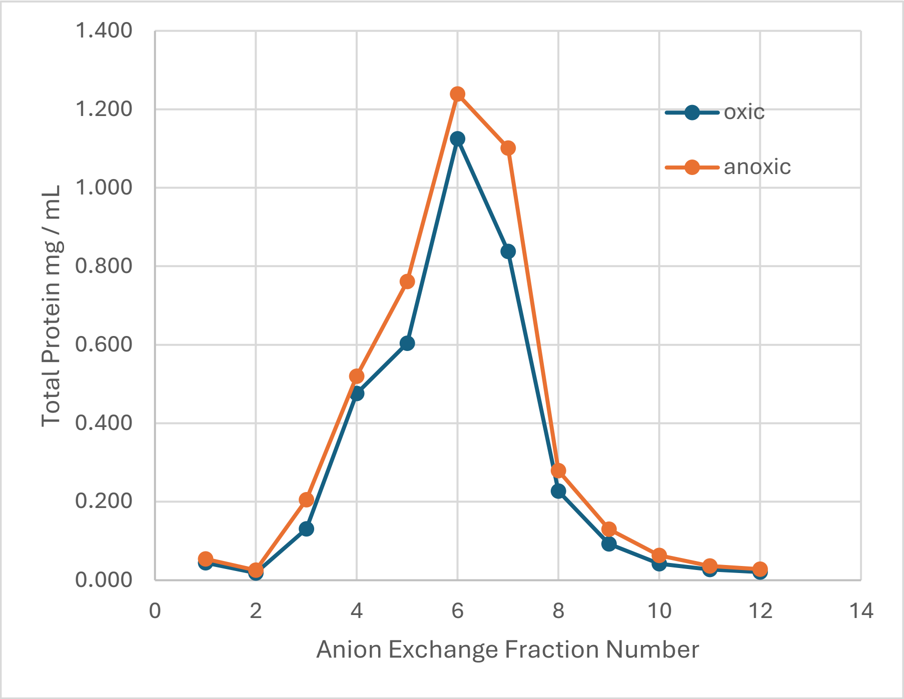


**Figure S1.** Total protein concentration data on the anion exchange fractions. See Table S1 for more information.

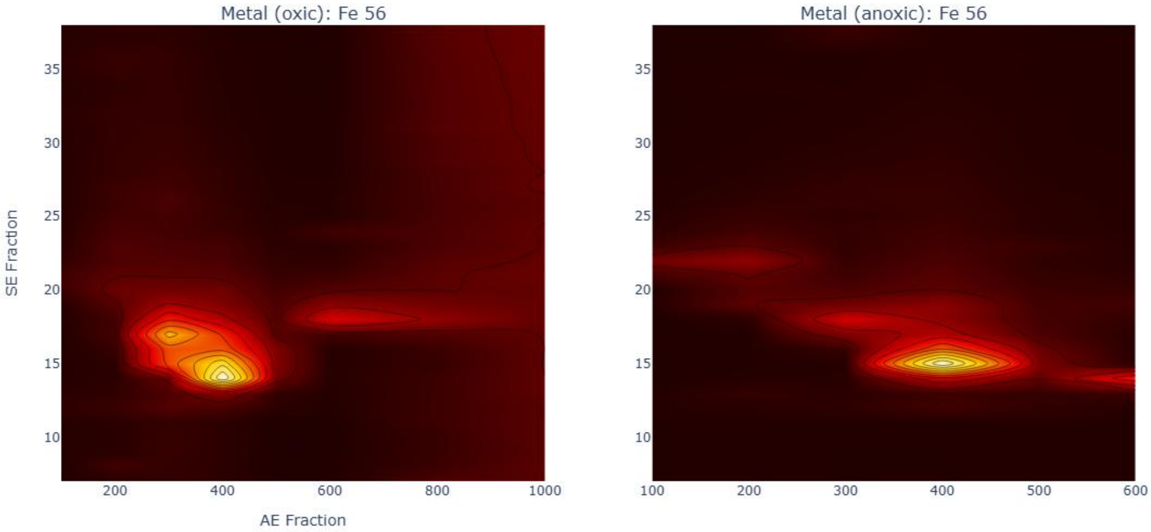


**Figure S2.** Enlarged 2d representation of iron metallome with extended range for oxic fraction (AE100-1000) for oxic. Similar to Figure 2 but without inset Fe Peak numbers.

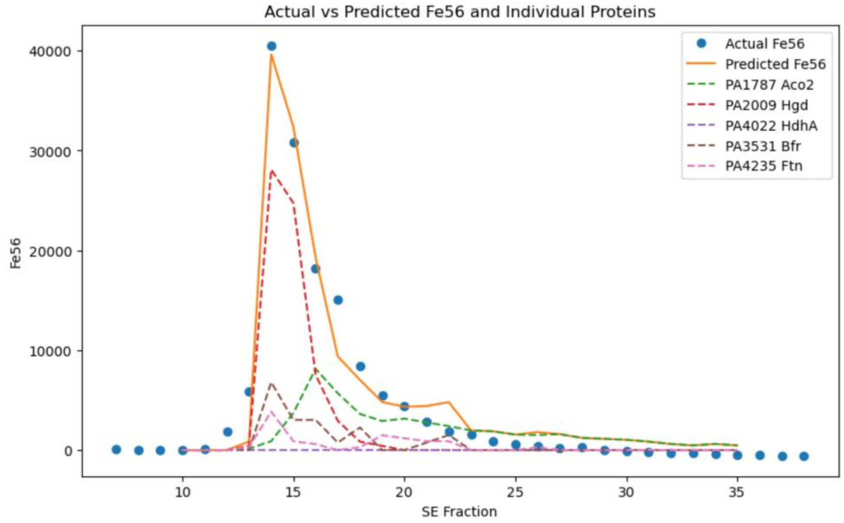


**Figure S3.** When the second peak of PA2009 is removed at SE 23 (AE 400), PA2009 contributes 45% to Peak 1.

**Figure S4.** CLUSTAL O(1.2.4) multiple sequence alignment of two *P. aeruginosa* bacterioferritin PA3531 and ferritin PA4235. The full amino acid sequences were verified to have no shared tryptic peptides. The alignment was generated at EBI's Clustal Omega portal https://www.ebi.ac.uk/jdispatcher/msa/clustalo.

```
PA3531          MKGDKKVIQHLNKILGNELIAINQYFLHSRMWNDWGLKRLGAHEYHESIDEMKHADKLIE  60
PA4235          MQGHPEVIDYLNTLLTGELAARDQYFIHSRMYEDWGFSKLYERLNHEMEEETQHADALLR  60
                *:*. :**::**.:* .** * :***:****::***:.:*  :  **  :* :*** *:.

PA3531          RILFLEGLPNLQDLGKLLIGENTQEMLQCDLNLELKATKDLREAIVHCEQVHDYVSRDLL  120
PA4235          RILLLEGTPRMRP-DDIHPGTTVPEMLEADLKLERHVRAALAKGIALCEQHKDFVSRDIL  119
                ***:*** *.::  ..:  * .. ***:.**:** :.   * :.*. *** :*:****:*

PA3531          KDILE-SEEEHIDYLETQLGLIQKVGLENYLQSHMHEDD  158
PA4235          KAQLADTEEDHAYWLEQQLGLIARMGLENYLQSQI----  154
                *  *  :**:*  :** ***** ::********::
```

**Figure S5.** CLUSTAL O(1.2.4) multiple sequence alignment of three *P. aeruginosa* ferritins, two bacterioferritins PA3531 and PA4235 as in Figure S4, and adding ferritin PA4235. The three full amino acid sequences were verified to have no shared tryptic peptides. The alignment was generated at EBI's Clustal Omega portal https://www.ebi.ac.uk/jdispatcher/msa/clustalo.

```
PA4880          MTTVQLTDVQTLRDRARKNIQEGAVTEGYSADRQTVLRLLNEALATELVCFLRYKRHYFM  60
PA3531          ------------------------------MKGDKKVIQHLNKILGNELIAINQYFLHSRM  31
PA4235          ------------------------------MQGHPEVIDYLNTLLTGELAARDQYFIHSRM  31
                                               .   *:  **  *  ** .  :*  *  *

PA4880          ATGLKASIAAAEFLEHANQEMQHADQLAERIMQLGGEPDFNPRGLEERSHAEYVEGKTLK  120
PA3531          WNDWGLKRLGAHEYHESIDEMKHADKLIERILFLEGLPNLQDLGK-------LLIGENTQ  84
PA4235          YEDWGFSKLYERLNHEMEEETQHADALLRRILLLEGTPRMRP-DD-------IHPGTTVP  83
                  .   .    .  ..  :* :*** * .**: * * * :.  .           * .

PA4880          DMVTENLIAERIAIDSYREIIT-YLGNDDPTTRRIFEEILAQ-EEEHADDMADILDDLA-  177
PA3531          EMLQCDLNLELKATKDLREAIVHCEQVHDYVSRDLLKDILE-SEEEHIDYLETQLGLIQK  143
PA4235          EMLEADLKLERHVRAALAKGIALCEQHKDFVSRDILKAQLADTEEDHAYWLEQQLGLIAR  143
                :*:  :*  *  .     : *.     .* .:* :::  *   **:*   :   *. :

PA4880          ---------------      177
PA3531          VGLENYLQSHMHEDD      158
PA4235          MGLENYLQSQI----      154
```

**Figure S6**. CLUSTAL O(1.2.4) multiple sequence alignment of two *P. aeruginosa* bacterioferritins PA3531 and PA4880. The full amino acid sequences were verified to have no shared tryptic peptides. The alignment was generated at EBI's Clustal Omega portal https://www.ebi.ac.uk/jdispatcher/msa/clustalo.

```
PA3531          ------------------------------MKGDKKVIQHLNKILGNELIAINQYFLHSRM  31
PA4880          MTTVQLTDVQTLRDRARKNIQEGAVTEGYSADRQTVLRLLNEALATELVCFLRYKRHYFM  60
                                               . :.*:: **: *..**:.: :*  *  *

PA3531          WNDWGLKRLGAHEYHESIDEMKHADKLIERILFLEGLPNLQDLGKLL-------IGENTQ  84
PA4880          ATGLKASIAAAEFLEHANQEMQHADQLAERIMQLGGEPDFNPRGLEERSHAEYVEGKTLK  120
                 ..   .  .*.  ..: :**:***:* ***: * * *:::  *            *:. :

PA3531          EMLQCDLNLELKATKDLREAIVHCEQVHDYVSRDLLKDILESEEEHIDYLETQLGLIQKV  144
PA4880          DMVTENLIAERIAIDSYREI-ITYLGNDDPTTRRIFEEILAQEEEHADDMADILDDLA--  177
                :*:  :*  *  * .. **  :     .* .:* ::::** .**** * :   *. :
```

```
PA3531      GLENYLQSHMHEDD 158
PA4880      -------------- 177
```
